# Multi-omics, organoid-based modeling reveals an SRC/mTOR-dependent fetal-like stem cell trajectory in colorectal cancer

**DOI:** 10.64898/2026.07.03.735750

**Authors:** Theresa Mulholland, Bogac Aybey, Zhenchong Li, Luisa Schwarzmüller, Niklas Rindtorff, Lucrezia Tondo, Ping Sui, Ekin Karabati, Philipp Albrecht, Julian E. Riedesser, Yvonne Petersen, Thilo Miersch, Erica Valentini, Elke Burgermeister, Tianzuo Zhan, Lena Dreikhausen, Nadine Schulte, Sebastian Belle, Stefan Wiemann, Michael Boutros, Matthias P. Ebert, Johannes Betge

## Abstract

**Background:** Single-cell atlases have described diverse stem cell states in colorectal cancer (CRC), however, the overarching trajectories of those states and the underlying functional mechanisms, including their relevance for drug sensitivity, need better understanding.

**Methods:** We established 64 patient-derived organoids from microsatellite-stable colorectal cancers, characterized their transcriptomes and genomes, and performed drug screening with 62-140 clinically approved substances. We analyzed additional published transcriptome data from patient-derived organoids (72 patients from three independent datasets), TCGA-CRC data (466 patients), and single-cell transcriptomes of tumor biopsies (123,000 cells from six independent cohorts) to establish a functional and molecular landscape of CRC stem cells. We performed mechanistic follow-up analyses by mass-spectrometry-based proteomics, large-scale kinase inhibition assays and immunofluorescence analyses.

**Results:** We find a continuous landscape of CRC stem cells that is characterized by distinct developmental programs: adult stem cell-to fetal-like regenerative states and transition between differentiation programs. By large-scale drug perturbations and multi-omics modeling, we identify a regenerative/fetal-like stem cell trajectory characterized by PI3K/mTOR dependency. We find the identified developmental axes conserved in organoid, clinical, as well as single-cell data, and the fetal-like PI3K/mTOR-dependent state to be associated with poor clinical prognosis. Mechanistically, PI3K/mTOR vulnerability is linked to a lack of adaptive capability due to suppressed mRNA translation and associated with an upregulated SRC signaling network.

**Conclusions:** Our work moves beyond a molecular CRC landscape by combined functional perturbation analyses in organoids. This enables mechanistic modeling of stem cell state regulation and identifies an SRC/mTOR-dependent regenerative state in CRC, which might allow improved therapeutic targeting in the future.

## Introduction

Colorectal cancer (CRC) is the third most common cancer worldwide and second leading cause of cancer-related deaths [1]. Prognosis is particularly poor in metastatic CRC, for which chemotherapy is still the backbone of treatment in the majority of cases. Although molecular stratification has enabled targeted therapies for specific patient subsets, most patients lack robust predictive biomarkers or novel targeting strategies to guide treatment selection. This reflects the substantial inter-and intratumoral heterogeneity of CRC.

Transcriptional states play an essential role in CRC drug sensitivity. Existing genome-and transcriptome-based CRC tumor classification systems [2–4], such as the Consensus Molecular Subtypes (CMS) [2], have provided important insights into CRC biology by defining discrete tumor categories. However, discrete CRC subtypes incompletely capture dynamic processes and have limited translational value for therapy stratification. Recent advances in developmental and single-cell biology have shown that epithelial identity is inherently dynamic [5–8]. CRC cells can transition between differentiated lineages, including absorptive and secretory programs, and adopt distinct stem-like states depending on context. Notably, for the latter, both adult intestinal stem cell programs and fetal or regenerative states have been implicated in tumor progression [9,10], metastasis [8], and therapy resistance [11–14]. How these programs are organized in human CRC, and whether they reflect distinct or overlapping biological trajectories, remains unclear. Furthermore, epithelial stem cell states and their relevance for drug sensitivity are still insufficiently characterized in CRC.

Here, we hypothesized that CRC epithelial programs are organized along continuous transcriptional axes rather than discrete subtypes, and that these axes are functionally linked to drug sensitivity. To address this, we leveraged patient-derived organoids (PDOs), which represent the molecular diversity of human CRC and preserve intrinsic epithelial programs, while enabling systematic multi-omics profiling and drug perturbation screening. By integrating gene expression, mutation status, and drug response across multiple datasets, we define a conserved epithelial landscape of CRC. We show that this landscape is structured by distinct developmental programs, separating fetal-like regenerative states from proliferative epithelial programs and metabolic programs from plasticity-related states. Importantly, these axes are directly associated with drug vulnerabilities, including differential sensitivity to PI3K/mTOR pathway inhibition associated with a plasticity-related fetal-like stem cell state. We find that this PI3K/mTOR vulnerability is linked to an inability to adapt to PI3K/mTOR inhibition due to suppressed mRNA translation and driven by an upregulated SRC signaling network, providing a framework to connect epithelial plasticity with underlying signaling nodes.

## Results

### CRC PDOs define two continuous axes of epithelial plasticity

To systematically investigate how tumor-intrinsic epithelial plasticity relates to drug vulnerabilities, we established a CRC PDO biobank from microsatellite stable tumors (MSS), comprising 132 samples derived from 64 individual patients, with several biological replicates available for a most patients, and performed transcriptomic and genomic characterization and functional drug screening of this cohort (Fig. 1a). To capture a wide range of heterogeneity, we included clinically diverse patients (Supp. Tab. 1, Supp. Fig. 1a). Our cohort mainly consists of treatment-naïve patients (n=58) from rectal (n=43) or colon (n=21) adenocarcinomas mostly from stage III-IV (n=42). Most prevalent mutations were *APC*, *TP53*, and *KRAS* (81%, 67%, and 65% of our cohort, respectively). This design allowed us to capture genomic variability across a broad spectrum of tumor epithelial subtypes and states.

**Fig 1.**
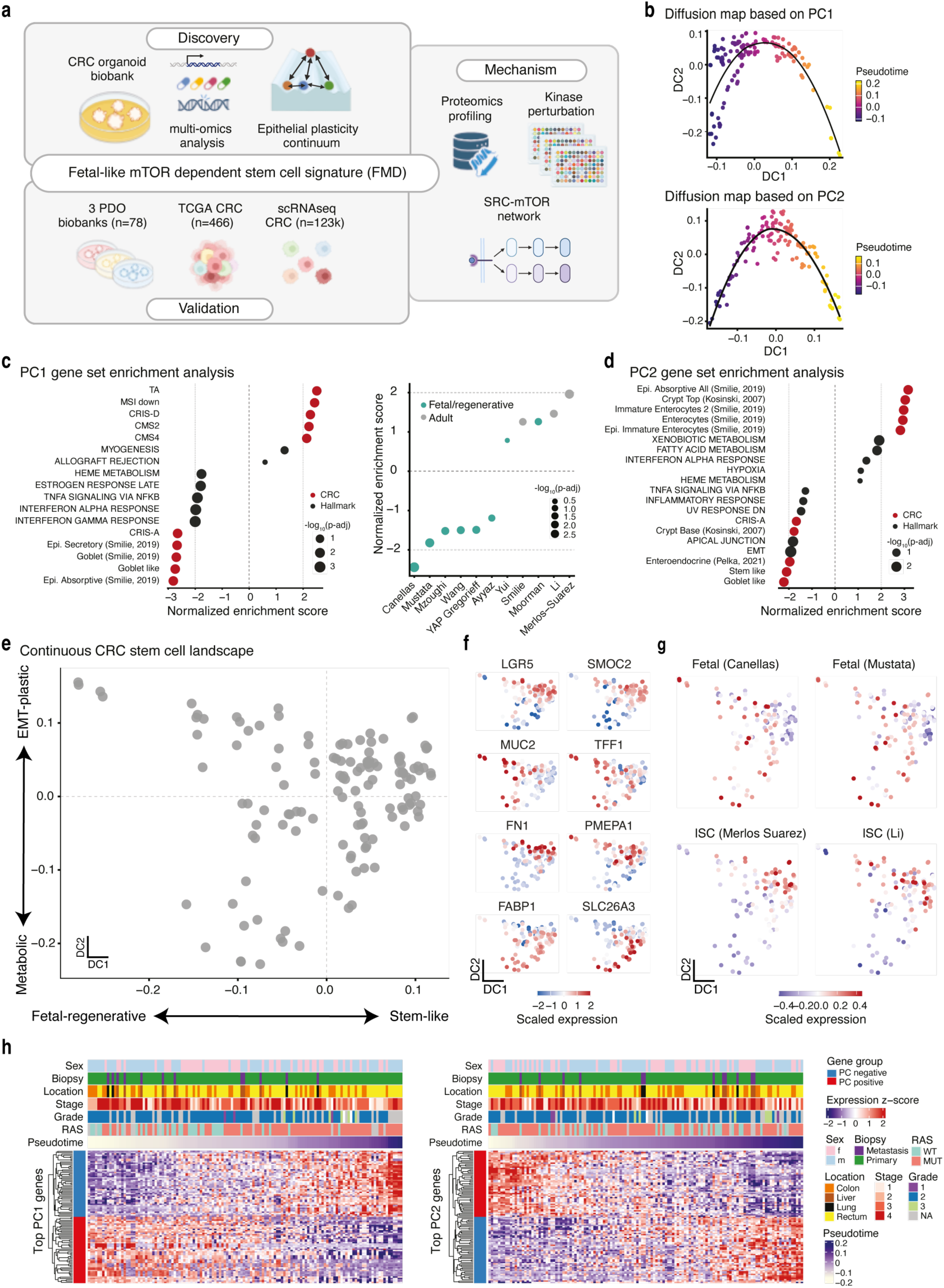
A continuum of epithelial plasticity in CRC PDOs. **(a)** Overview of experiment and analysis design. **(b)** Diffusion maps based on top 50 PC1 or PC2 probes. For the diffusion maps, probes with the 50 highest and 50 lowest loading values from PC1 (left) or PC2 (right) are used. Pseudotime values are inferred from the diffusion map, showing lower (dark magenta) to higher pseudotime (yellow). Samples align along continuous trajectories. Gene set enrichment analysis for **(c)** PC1 or **(d)** PC2 axes. Genes are ranked by loading values. CRC specific, intestinal stem cell (ISC), fetal/regenerative and YAP signatures are used and marked by colors. Normalized enrichment scores are shown along with their corresponding FDR-adjusted p-values in bubbles. Bubble sizes show-log_10_(FDR) values. **(e)** Joint diffusion map representing the CRC stem cell plasticity landscape. Diffusion map is constructed based on mutually exclusive highest and lowest top 50 PC1 and PC2 probes (n(PC1+, PC1-, PC2+, PC2-): 47, 46, 47, 46, respectively). Samples align along two continuous axes: metabolic vs. EMT-plastic and intestinal stem-like vs. fetal/regenerative. **(f)** Marker gene expression on joint diffusion map. Z-scaled marker gene expression values for selected absorptive/metabolic, EMT/mesenchymal/plastic, secretory, and stem-like programs are shown along the joint diffusion map. Markers align across distinct biological gradients. **(g)** Fetal and intestinal stem cell signature scores on joint diffusion map. Mean signature scores for two ISC and two fetal/regenerative signatures are shown along the joint diffusion map. The first diffusion component separates regenerative/fetal vs. intestinal stem-like programs. **(h)** Continuous expression of top 50 PC1 or PC2 probes. Z-scaled expression values for each top PC1 (left) and PC2 (right) probes are depicted as heatmap. Expression values are shown from low (blue) to high (red). Both sides of PC probes are colored separately: positive (red) and negative (blue) side. Samples are ordered along the pseudotime inferred from diffusion maps and gene expression patterns show continuous gradients.

First, we aimed to define a landscape of epithelial stem cell states in CRC based on gene expression analyses of our PDOs. Gender-adjusted PCA did not show discrete clusters, rather a continuous alignment of samples across first and second principal components (PC1 and PC2) (Supp. Fig. 1b). We further examined if dominant variations captured by PC1 and PC2 reflect continuous gradients. We constructed diffusion maps using the probes with the highest and lowest PC loadings (50 each, denoted as PC+ or PC-, respectively) for PC1 and PC2, separately (Fig. 1b, Supp. Tab. 2 for probe lists). The selection of 50 probes was a pragmatic feature-selection approach to capture the dominant transcriptional programs represented by each PC while limiting noise from lower-loading probes. This yielded focused and biologically interpretable probe sets. Samples were ordered along single continuous trajectories in both PC1 and PC2-based diffusion maps. Heatmaps visualize these gradual expression changes of the PC-derived probes across the resulting pseudotime ordering (Fig. 1h). To assess whether these trajectories reflected coherent biological structure rather than arbitrary probe selection, we compared the resulting embeddings to size-matched random probe sets. PC-derived probe sets showed stronger alignment with pseudotime and the first diffusion component than random probe sets, showing that the observed trajectories were not driven by random probe selection but reflect a coherent ordering captured by the top PC probes (Methods, Supp. Fig. 1c). These initial results showed that CRC PDO samples distribute along continuous axes rather than randomly or forming discrete clusters.

Then, we characterized PC1 and PC2 axes separately to examine the underlying epithelial biology. The PC1+ axis was enriched for cellular development, and transit-amplifying as well as intestinal stem cell signatures (Fig. 1c, Supp. Fig. 2a), supported by canonical stem and developmental markers, such as *LGR5*, *HMGA2*, and *IGF2*. The PC1-axis was enriched for interferon, TGFβ, secretion, inflammation, goblet-like, YAP and fetal/regenerative signatures, with representative genes including *MUC2*, *TFF1*, and *REG4* (Fig. 1c, Supp. Fig. 2b). Together, PC1 captures a continuum between fetal/regenerative and intestinal stem-like programs.

The PC2+ axis was enriched for absorptive and metabolic pathways, supported by enterocyte-associated genes such as *ACE2*, *DPP4*, and *FABP1* (Fig. 1d, Supp. Fig. 2b, Supp. Fig. 2d). In contrast, the PC2-axis was enriched for pathways linked to epithelial modeling and EMT-associated processes, accompanied by increased expression of genes involved in extracellular matrix organization and cellular migration, including *FN1*, *CXCR4*, and *PMEPA1*. Rather than reflecting a fully mesenchymal state, this axis was consistent with epithelial plasticity and partial EMT-like programs. Accordingly, PC2 defined an orthogonal dimension of epithelial heterogeneity that separates differentiated metabolic states from remodeling-associated, EMT-like cellular programs.

To assess the relationship between our plasticity axes and established colorectal cancer classification systems, we quantified gene-level overlap between the top 50 PC-derived probe sets and published CMS [2], CRIS[3], and iCMS [4] signatures. Direct overlap was limited (Jaccard indices generally < 0.1) indicating little global redundancy with existing classifier genes. However, in our data, PC1 score correlated positively with CMS2, iCMS2, and CRISD scores (r = 0.6), and negatively with iCMS3, CMS1/CMS3, and CRISA/CRISB scores (r =-0.7 to-0.8). PC2 scores showed selective positive correlations with CMS2 and CRISC signatures (r=0.6) and negative correlations with CMS1 and CRISA signatures (r =-0.3). Overall, these axes partially intersect with known subtype programs while capturing complementary transcriptional variation beyond established CRC classifiers.

Next, we integrated both axes together to create a unified framework of the epithelial plasticity landscape (Fig. 1e). To this end, we constructed a diffusion map based on the expression values of the mutually exclusive probes from top 50 PC1+/-and PC2+/-probes (n(PC1+, PC1-, PC2+, PC2-): 47, 46, 47, 46, respectively), thereby capturing the principal and orthogonal sources of transcriptional variation. This approach captures sample-to-sample similarities across both principal and orthogonal transcriptional programs within a shared continuous low-dimensional manifold. Within this embedding, representative marker genes for absorptive/metabolic, EMT-associated plasticity, secretory and stem-like programs aligned with our corresponding epithelial axes (Fig. 1e). In addition, fetal/regenerative and intestinal stemness signatures marked distinct ends of DC1 along our landscape (Fig. 1g). These analyses confirmed the biological interpretability of the projection and created our continuous CRC stem cell states landscape.

To assess whether clinical or genomic features define parts of the epithelial landscape, we projected main features onto the diffusion embedding (Supp. Fig. 3). Neither clinical data nor common driver mutations (*KRAS*/*NRAS*, *APC*, *TP53*, and *FBXW7*) formed discrete clusters, indicating that the epithelial trajectories are not primarily structured by genotype or sample origin. Nevertheless, activating mutations in the RAS pathway (*KRAS*/*NRAS*) were associated with a shift along the primary diffusion axis (DC1; linear model, *p* < 0.001), while showing no significant association with the orthogonal axis (DC2; *p* = 0.22). Notably, the variance explained by *KRAS*/*NRAS* status was modest (DC1, R² = 0.15), indicating that these mutations bias cellular positioning along the trajectory without defining its overall structure. Of note, such moderate association between *KRAS/NRAS* and inflammatory/fetal-regenerative programs have been already shown [10,11]. Further, anatomical location of PDO (left colon, right colon, or rectum) did not explain a significant amount of variance in either DC1 (R² =-0.02, *p* = 0.87) or DC2 (R² = 0.017, *p* = 0.17). Together *KRAS*/*RAS* status and location of tumor explained only 15% of variance in DC1 (*p* = 0.0013) and almost none in DC2 (R² =-0.003, *p* = 0.43). These results suggest that the epithelial plasticity landscape is predominantly governed by transcriptional state rather than discrete genomic alterations, with oncogenic mutations exerting a secondary, modulatory effect.

Together, we defined a continuous CRC epithelial plasticity manifold, structured by two independent and orthogonal biological axes that represent different stemness (adult vs. fetal-like/regenerative) and metabolic vs. plasticity related states.

### Multi-omics modeling identifies a PI3K/mTOR-dependent state within the epithelial plasticity landscape

Next, we sought to obtain gene expression modules explaining distinct drug vulnerabilities and to assess the link to our epithelial plasticity landscape. To represent multiple modalities such as transcriptomics, genomics and drug screening data, we built a Multi-Omics Factor Analysis (MOFA) model [15]. To increase the robustness and generalizability of our analysis, we used a discovery and validation strategy by splitting up our organoid drug library cohort into two, CRC1 and CRC2, restricting our initial analysis to the largest cohort with drug-PDO combinations (CRC1, 52 drugs-42 PDOs), which cover most parts of our epithelial landscape (Supp. Fig. 4). For the validation of our MOFA results, we included a second cohort, CRC2 (22 PDOs and 140 drugs), whose samples spanned similar regions along the stem cell plasticity axes. For the MOFA model, we used most prevalent mutations (mutated in at least three donors, n=12), gene expression values of top 10% highly variable genes (n=3812), and area under dose-response curves (AUC) values from 52 drugs. This framework enabled us to examine drug-transcriptome-genomics axes in a fine-granular and robust way.

MOFA analysis converged on six factors with distinct associations, cumulatively explaining 67 % of the variance in the data (Fig. 3a,b). The first latent factor was the one explaining the largest variance, cumulatively 20 %, in the dataset, contributing to the response of several drugs (16% of drug variance, Supp. Fig. 5) and was weakly associated with gene expression (2%) as well as mutations (2%). Factor 1 showed a high association with *RAS* mutation status and EGFR inhibitor resistance, which is an extensively studied drug-genotype association [16] (Supp. Fig. 5).

The most informative latent factor in the context of drug-gene expression associations was the second factor (F2): explaining both variance in gene expression (6%) and drug response (11%). The top drug loadings of Factor 2 were PI3K/mTOR-pathway inhibiting drugs including Vistusertib, Gedatolisib, Taselisib, and Everolimus. It also included inhibitors of the RAS/MAPK signaling pathway (MEK inhibitors, BRAF inhibitors), known to interact with the PI3K/mTOR pathway, as well as multi-tyrosine kinase inhibitors and a CBP/beta-catenin inhibitor (Fig. 2b-c). Similarly, AUCs between Factor 2 high vs. low organoids showed marked differences (Fig. 2d). Of note, compounds targeting PI3K/mTOR were also among the top drug loadings of Factor 1, but overall target association to Factor 1 was less than in Factor 2 (Fig. 2b).

**Fig 2.**
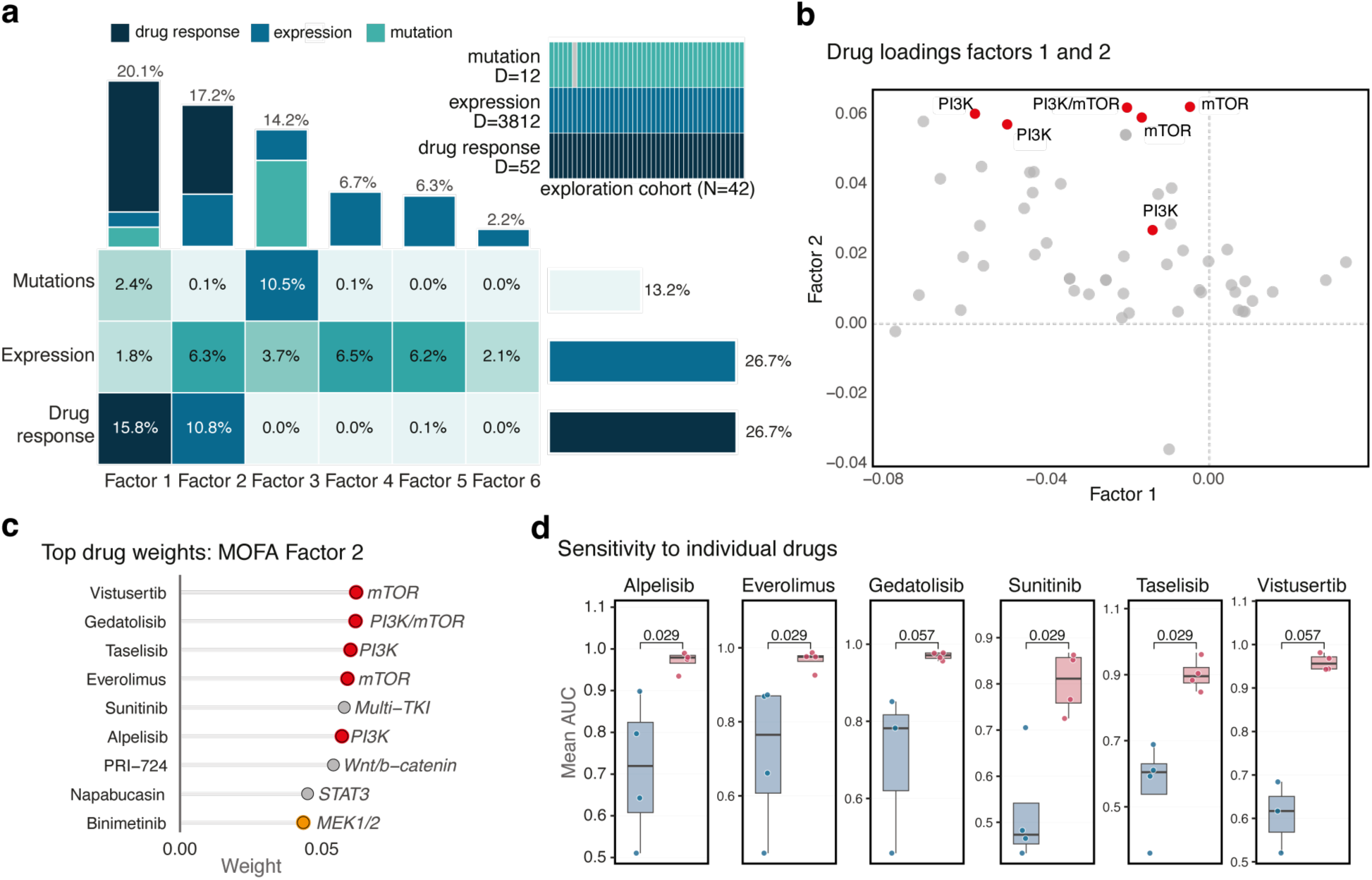
MOFA identifies latent axes linking drug response, gene expression, and mutation status in CRC organoids. **(a)** MOFA analysis between drug response, gene expression and mutation status in CRC cohort 1 (exploration cohort). As input for MOFA, 10% most variable probes (n=3812), DMSO-normalized AUC values for 52 drugs and genes (n=12) with mutation in at least three samples are used. Six distinct latent factors are obtained. Variance explained by latent factors are shown from low (white/green) to high (dark blue). **(b)** Drug loadings of factors 1 and 2. PI3K/mTOR targeting drugs are highlighted **(c)** Drugs with the highest associations with MOFA Factor 2. Factor weights of top drugs are depicted. **(d)** Response of organoids with highest vs. lowest Factor 2 values to selected drugs with highest factor 2 loadings. All p-values are calculated using Wilcoxon rank-sum test (two-sided).

**Fig 3.**
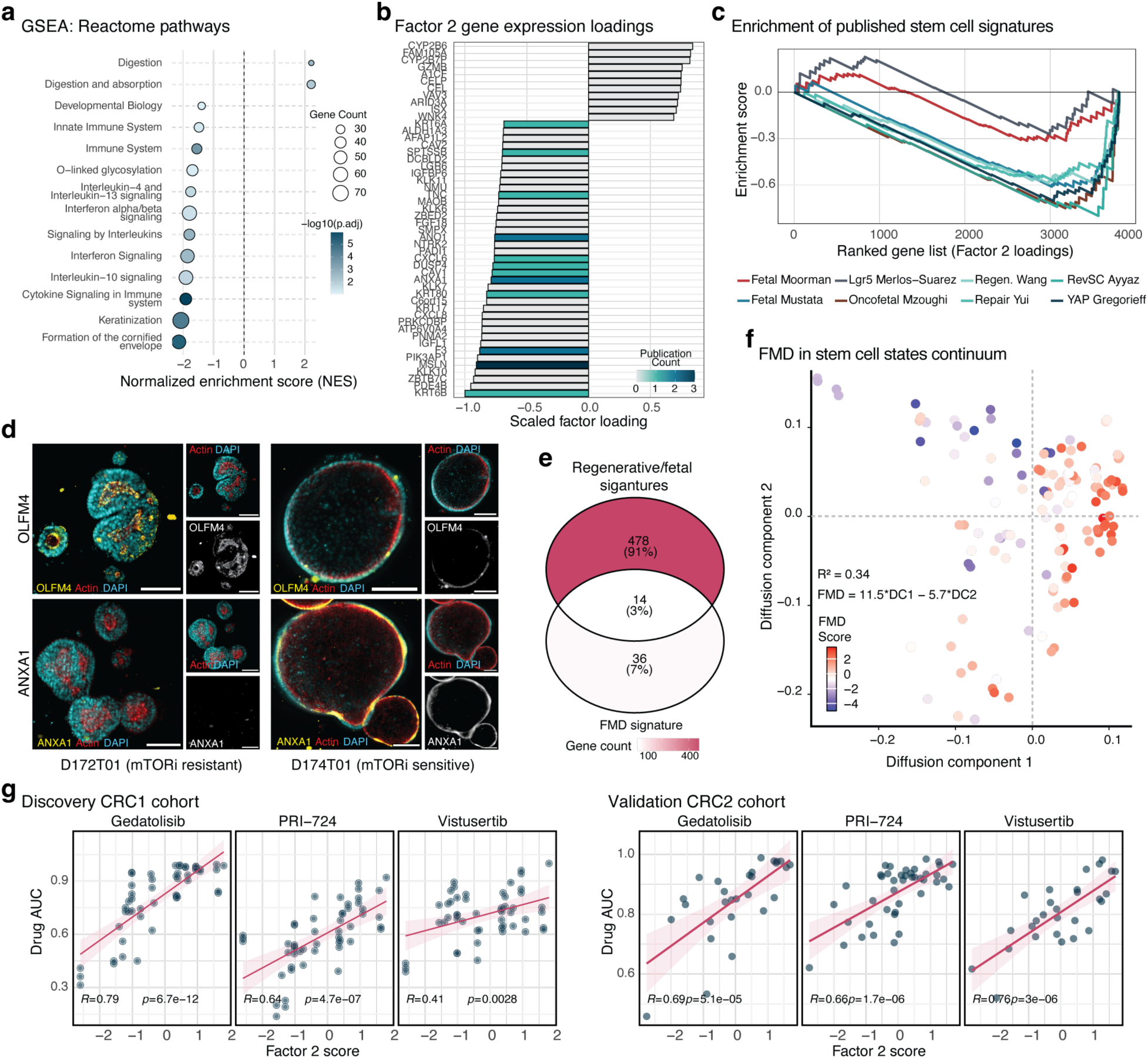
**An PI3K/mTOR dependency signature is associated with epithelial plasticity programs**. **(a)** Gene set enrichment analysis of Factor 2 loadings. Genes are ranked by their Factor 2 weights (Supp. Table 5). Normalized enrichment scores are calculated for Reactome pathway signatures. **(b)** Top genes associated with MOFA Factor 2. Z-scaled factor loading values are shown for the highest-ranking genes. Genes previously associated with fetal or regenerative stem cell signatures are colored, signatures published by Mustata et al. (2013) [17], Gregorieff et al. (2015) [18], Yui et al. (2018) [19], Ayyaz et al. (2019) [20], Wang et al. (2019) [7], Moorman et al. (2025) [8], Merlos-Suárez et al. (2011) [21] and Mzoughi et al.(2025) [10] are considered. **(c)** Enrichment scores for fetal/regenerative and intestinal stem cell signatures across Factor 2 loadings. Genes are ranked by their Factor 2 weights and enrichment scores are computed for each signature gene. **(d)** Representative images of two PDO lines (mTOR-resistant D172, mTOR-sensitive D174) showing ANXA1 and OLFM4 and DAPI. Split channels are shown to illustrate marker-specific staining patterns. Scale bars 100µm. **(e)** Overlap of the fetal-mTOR-dependency gene set (FMD gene set) with previously published fetal/regenerative signatures. **(f)** Mapping of the FMD signature across CRC stem cell plasticity axes. FMD score is defined as the mean z-scaled expression of the top 50 positively loaded genes minus the mean z-scaled expression of the top 50 negatively loaded genes. FMD score is regressed against DC1 and DC2, with R² values indicating the proportion of variance explained. Higher and lower FMD scores are shown in red and blue, respectively. **(g)** Association between AUC values and FMD score for three drugs with high association with factor 2 in the discovery cohort and validation cohort. The correlation between FMD score and AUC values is shown using Pearson correlation.

We then characterized F2 and derived PI3K/mTOR sensitivity-associated gene modules by integrating its associations with gene expression, drug sensitivity, and epithelial plasticity axes. Samples with low F2 loadings, corresponding to mTOR-sensitive PDOs, were enriched for fetal/regenerative programs, including inflammatory, interferon, and immune system activation, whereas resistant samples showed increased activation of absorptive and metabolic programs (Fig. 3a-c). Notably, comparison with published fetal signatures revealed only partial overlap among the top F2-associated genes (36 unique and 14 shared genes, Fig. 3E), suggesting that the fetal-mTOR-dependent program represented by F2 captures both established fetal-like programs and additional fetal-state and mTOR-dependency genes not previously linked to stemness programs (Supp. Fig. 6) These suggested that the fetal-mTOR dependent program represented by F2 captures both established fetal-like programs and further fetal state genes and/or mTOR dependency genes which have not been linked to stemness programs. Consistent with this interpretation, immunofluorescence staining of two representative PDOs showed increased expression of ANXA1 (fetal marker and top loaded gene in Factor 2) and a lack of OLFM4 expression (marker of LGR5+ stem cells) in mTOR-sensitive PDOs compared to resistant PDOs (Fig. 3d). To quantify this mTOR-dependency axis, we defined an mTOR sensitivity score (fetal-mTOR-dependency score, FMD score) as the difference between the average expression of the 50 top and bottom-ranked genes based on F2 loadings. In both organoid cohorts, the FMD score showed strong correlation with mTOR and CBP/beta-catenin inhibitor response (AUC; Pearson r = 0.7-0.8; Fig. 3g left), and this relationship was consistently reproduced in the independent drug library cohort (Pearson r = 0.4-0.8; Fig. 3g right), demonstrating robustness across datasets. Among all factors, FMD showed the strongest association with the epithelial axes, correlating with both PC1 and PC2 directions (Pearson r(F2+, PC1+| PC2+) and r(F2-, PC1-| PC2-) > 0.5), thereby linking mTOR sensitivity to a specific combination of epithelial plasticity programs (Supp. Fig. 7). Further, mapping these scores onto the epithelial plasticity landscape revealed that regions enriched for mTOR dependency showed higher EMT-plastic and fetal/regenerative programs and lower metabolic and adult-like programs (Fig. 3f).

Of note, the third MOFA factor was associated with *RAS* mutation status and metabolism-related gene expression (negative factor loadings, Supp. Fig. 8). Other factors (factor 4-6) solely accounted for variance in gene expression data and were not of primary interest for further analyses.

In conclusion, integrating our CRC stem cell landscape with a multi-omics modeling approach allowed us to retrieve a gene expression signature for PI3K/mTOR dependency and associate this with a subset of samples on the fetal/regenerative stem cell trajectory.

### Plasticity axes and PI3K/mTOR-dependency signature are conserved across PDOs and patient tumors

Next, we asked how FMD signature aligns with the CRC stem cell plasticity axis across different CRC datasets. To this end, we included human MSS tumor expression datasets of different origins: PDO biobanks, patients’ bulk tumor samples from TCGA CRC (COAD/READ) and CRC biopsy samples profiled by single-cell RNA sequencing. This comprehensive evaluation aimed to establish a strong foundation for transfer of findings across platforms and modalities. For each dataset, we computed and represented the FMD score on individual plasticity axes in our cohorts as well as in the validation cohorts. Then, we showed the existence of our epithelial axes and its relation to the FMD score on the different datasets.

First, we were able to recapitulate our axes in external PDO datasets. To fully capture heterogeneity, we integrated RMA-normalized microarray expression data from three different PDO cohorts (n=78) [22–24]. On the diffusion map based on our axis genes (Fig. 4a), samples from independent cohorts occupied overlapping positions along the axis and did not primarily cluster by cohort (Supp. Fig. 9a). Importantly, the diffusion map based on our axis genes revealed the same epithelial state space based on the marker genes from our previous analysis, including the subset trajectory characterized by mTOR inhibitor sensitivity. Similarly, expression of our axis genes showed continuous gradients where samples from each cohort were aligned on the trajectories (Supp. Fig. 10a). This underlined that including multiple datasets was important to cover a higher level of heterogeneity not covered by individual datasets. These results indicated that our epithelial plasticity axes, including the FMD trajectory, can be reproduced in other independent PDO datasets.

**Fig 4.**
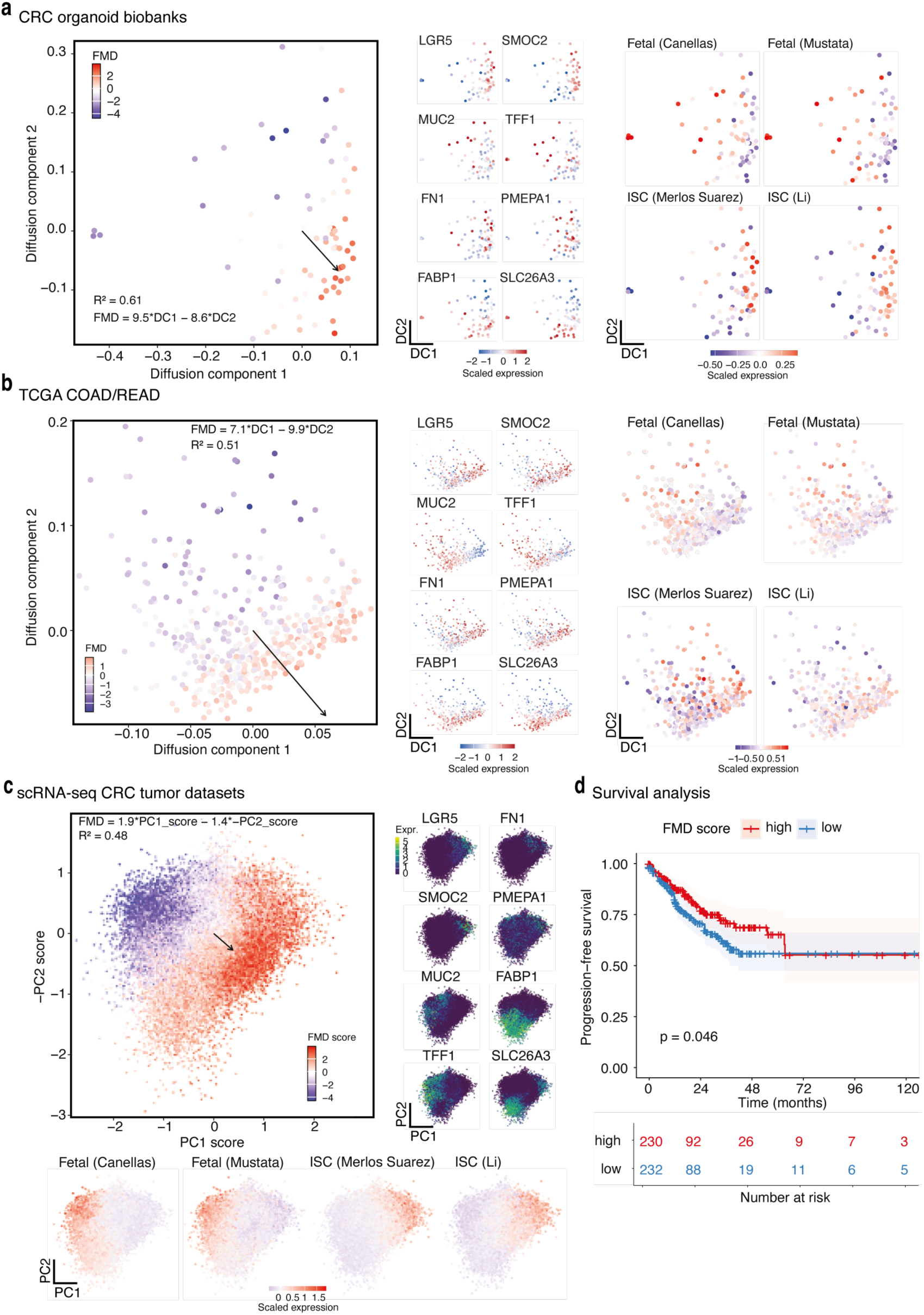
Epithelial plasticity axes and PI3K/mTOR-associated signature are conserved across independent CRC datasets. Representation of the fetal-mTOR-dependency (FMD) axis (MOFA Factor 2 transcriptional signature) across independent CRC datasets including **(a)** three CRC PDO biobanks, **(b)** TCGA CRC cohorts (COAD/READ) and **(c)** six single-cell CRC datasets. For PDO and TCGA datasets, diffusion maps are constructed using epithelial plasticity axis genes, and z-scaled expression values of representative marker genes are projected onto the embeddings from negative (blue, mTOR inhibitor sensitive / high fetal markers) to positive/high (red, mTOR inhibitor resistant, low fetal markers). For single-cell datasets, PC1 and PC2 epithelial program scores are calculated as the difference between the average expression of positive-and negative-end genes, and normalized expression values of representative marker genes are projected onto diffusion embeddings from low (dark magenta, mTOR inhibitor sensitive) to high (green-yellow, mTOR inhibitor resistant). Fetal/regenerative and intestinal stem cell signature scores are mapped across embeddings to assess conservation of epithelial plasticity programs. FMD scores are generally aligned with epithelial plasticity axes across datasets, supporting a conserved association between epithelial state and PI3K/mTOR response programs. **(d)** Association of the FMD signature with progression-free survival in TCGA CRC. Samples are stratified into FMD-high (mTOR inhibitor resistant / low fetal markers) and FMD-low (mTOR inhibitor sensitive, low fetal markers) groups using the median FMD score, and progression-free survival is compared by Kaplan–Meier analysis.

Our second validation model examined expression values (TPM-normalized) from patients’ bulk tissue CRC datasets from TCGA (COAD/READ, n=466). On the diffusion map, the samples formed the same gradients across two axes and two cohorts, COAD and READ, and did not form distinct clusters (Fig. 4b, Supp. Fig. 10b). At the single-gene level, despite the heterogeneity due to bulk tissue sequencing, PC gene modules remained coherent and formed the continuous gradient along individual trajectories (Supp. Fig. 11b). Expectedly, multiple programs were activated simultaneously in some samples due to the bulk tissue setting and those programs potentially being influenced by non-cancerous cells such as stromal cells [25]. Thus, even in a more heterogenous model, we could project bulk tissue samples onto our axes and demonstrate continuous gradient expression patterns along same epithelial plasticity landscape.

Within this framework, established CRC subtypes did not form discrete clusters but instead distributed continuously along the epithelial plasticity landscape (Supp. Fig. 11c). While CMS2 and CMS4 samples were enriched toward metabolic and EMT-associated regions, CMS1 and CMS3 aligned toward more fetal/regenerative states, consistent with overlapping transcriptional programs across subtype definitions. Notably, states typically associated with microsatellite instability samples (MSI) and immune infiltration were detectable even in MSS tumors. In addition, *KRAS*-mutant tumors were preferentially positioned toward the fetal/regenerative end of the landscape, consistent with our PDO analysis, and likely reflecting increased inflammatory signaling associated with these mutations. These further support the notion that these programs reflect shared biological states rather than strictly genotype-defined categories.

Clinical and genomic variables did not partition samples into discrete groups along the trajectory but instead showed modest, axis-specific effects (Supp. Fig. 10c). Linear modeling revealed that tumor site, histological subtype, and *KRAS* status collectively explained a limited proportion of variance in DC1 (R² = 0.26) and in DC2 (R² = 0.17). Effect size analysis further indicated that histology had the strongest influence on DC1 and DC2 (Cliff’s δ = 0.8 and-0.6, respectively), whereas tumor site and *KRAS* status showed only small or negligible effects (δ < 0.33). These confirmed our previous analysis in our PDO biobank that epithelial plasticity landscape is primarily governed by transcriptional state, with clinical and genomic features contributing only secondary, modulatory effects.

TCGA COAD/READ allowed us to test the association of progression-free survival (PFS) and FMD score. Samples stratified by median FMD scores (high or low) showed significant differences in PFS (log-rank p = 0.046, Fig. 4d), mTOR sensitivity (low FMD score, subset of fetal/regenerative stem cell axis) being associated with worse outcome. In contrast, the association with overall survival was not statistically significant (log-rank p = 0.58, Supp. Fig. 10d). Further, in univariate analysis, FMD score showed a significant association as a continuous variable with PFS (HR = 0.79, 95% CI = 0.63–0.98, p = 0.033, Supp. Fig. 10e), while multivariate Cox proportional hazards analysis adjusted for age, gender, and pathological stage showed a significant association as well (HR = 0.74, 95% CI = 0.55–0.99, p = 0.042, Supp. Fig. 10f). These indicate an independent association between lower FMD scores and increased risk of progression.

Finally, we investigated single-cell CRC malignant epithelial samples (n=123,000) from six datasets from four independent studies [26–29]. Here, we constructed program scores for PC1 and PC2 modules due to the sparsity and technical noise inherent to single-cell expression data: subtracting average expression values of our PC-genes from our PC+ genes separately for PC1 and PC2. On the PC1-versus-PC2 program scores plot, malignant cells aligned in two continuous axes in geometry as well as in expression instead of forming discrete clusters (Fig. 4c, Supp. Fig. 11a). Based on the expression of representative markers of our axes, epithelial biological programs were activated in distinct parts of the continua. Further, samples did not cluster by the dataset, meaning that our axes capture the epithelial biology rather than dataset-specific effects and we capture overarching epithelial states (Supp. Fig. 11b). Interestingly, within individual tumors, multiple epithelial states co-existed, however, with variability in their patient-specific compositions (Supp. Fig. 11c), reflecting intratumoral heterogeneity and plasticity of malignant epithelial cells. We demonstrated that, at single-cell resolution, malignant tumor cells occupied the same continuous two-dimensional plasticity space defined in PDOs.

The FMD gradient on the plasticity axes revealed consistent spatial patterns of mTOR sensitivity across cohorts. We modeled FMD score as a function of the underlying diffusion components or PC scores. Across datasets, FMD score was well approximated as a linear combination of these axes, defining a consistent direction within the PC1–PC2 space. Both the spatial distribution of mTOR inhibition sensitive regions and their mathematical representation were highly concordant across systems. These results showed that mTOR dependency does not align with a single epithelial identity, but with combinations of plasticity programs and the associated regions were conserved across experimental systems.

Hence, our extensive analysis of organoid, clinical and single cell datasets demonstrated that we could project the previously identified CRC stem cell developmental axes onto tumors at different granularities and validate that tumor cells characterized by the fetal-mTOR-dependency signature represent a subset of regenerative/fetal stem cell states, associated with adverse outcome.

### PI3K/mTOR dependency is linked to a lack of adaptive capability due to suppressed mRNA translation

To investigate the mechanistic foundation of the observed mTOR dependency, we utilized our MOFA-derived Factor 2 loadings to select a representative cohort of eight PDOs, representing the highest (n=4) and lowest (n=4) extremes of the sensitivity axis (Fig. 5a). For a comprehensive analysis we performed drug perturbation experiments with mTOR inhibitors (and SN38 as control) and generated RNA-seq, proteomic and phospho-proteomic data at baseline and after perturbation. Baseline characterization of these PDOs using RNA-seq and proteomics confirmed that the FMD signature was retained robustly across both modalities (Supp. Fig. 12). GSEA based on RNA-seq data confirmed that mTOR sensitivity (Fig. 5b) was significantly associated with epithelial developmental and inflammatory programs, as well as interferon signaling, aligning with previous analyses derived from microarray data (Fig. 3a). Additionally, the RNA-seq dataset identified additional sensitivity-associated pathways, i.e., characterized by an upregulation of the YAP-Hippo pathway and a downregulation of translational machinery components (Fig. 5b). When specifically examining PI3K and mTOR signaling, sensitivity correlated with increased transcriptomic enrichment of Hallmark PI3K/AKT/mTOR signatures [30], partly driven by the adapter protein PIK3AP1, one of the genes with the highest factor loadings and previously linked to PI3K/AKT pathway activation [31]. However, this transcriptional association did not translate into increased functional pathway activity. Instead, both proteomic GSEA and imputed kinase activity analyses revealed marked repression of the PI3K/mTOR signaling axis in mTOR-sensitive PDOs (Fig. 5c-d). Specifically, sensitive PDOs showed reduced kinase activities of MTOR, AKT1 as well as RPSKA1, suggesting a failure to translate the transcriptomic state into active signaling. By integrating transcriptomic and proteomic changes between the sensitivity groups we identified signaling components essential to translation, such as EIF4E and RPS6KA1, which showed distinct decoupling, characterized by high mRNA abundance but significantly depleted protein levels (Fig. 5e). In contrast, other PI3K/mTOR pathway components, specifically metabolic adapters, such as SLC2A1 and SFN showed coupled upregulation in the mTOR sensitive PDO group. The suppression of the translational machinery (Fig. 5b) provides a putative mechanism for the observed proteomic bottleneck, where PI3K/mTOR signaling in sensitive models is restricted by a failure in translational output. Further suppression of PI3K/mTOR signaling by pathway inhibitors may thus be a specific vulnerability in these tumor cells. Two drivers of this fetal-like mTOR-dependent state that were significantly upregulated on RNA level remained upregulated on the proteomic level in sensitive PDOs as well. The calcium-binding protein ANXA1 as well as the B-cell adapter PIK3AP1 may serve as biomarkers of the FMD state (Fig. 5f).

**Fig 5.**
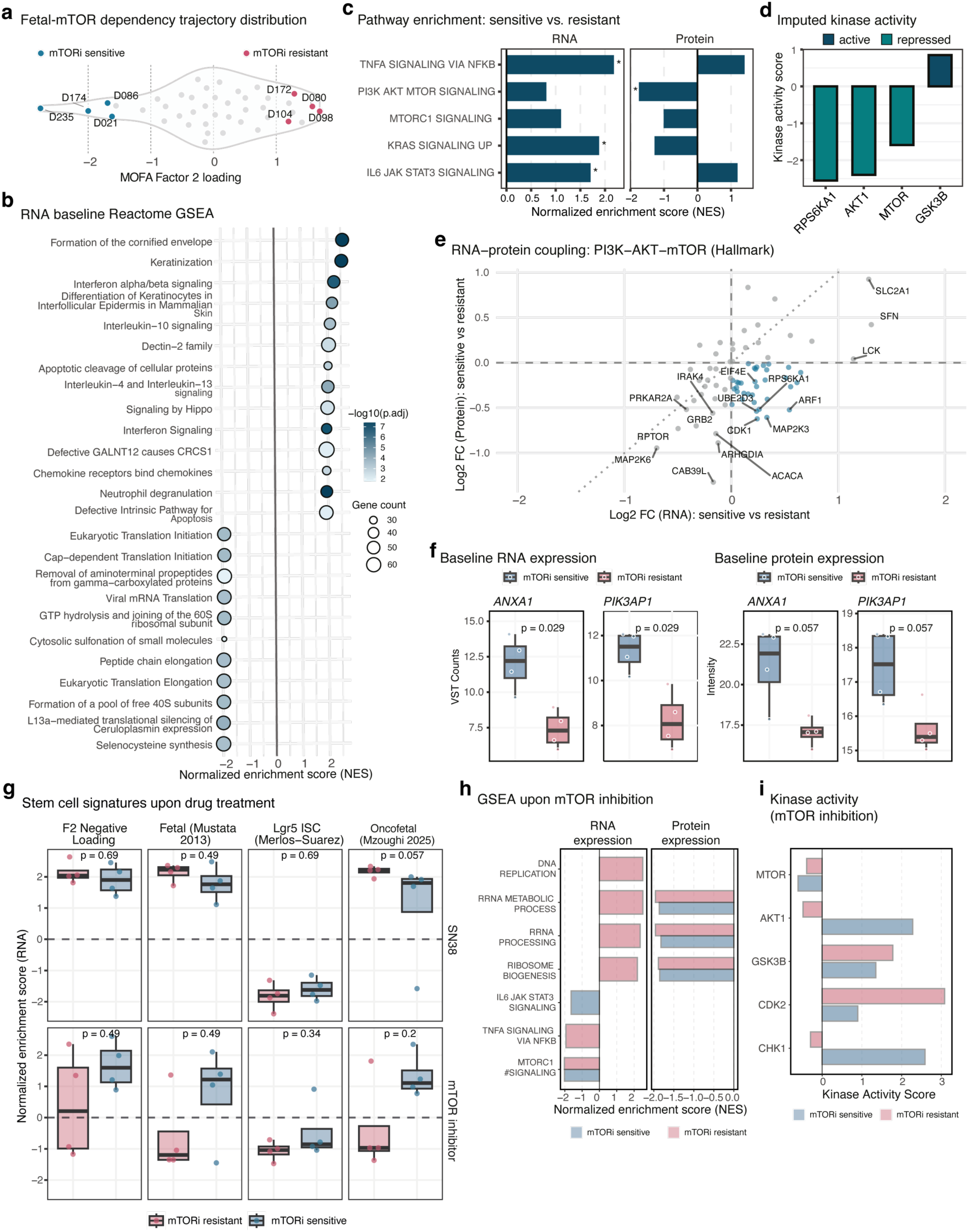
PI3K/mTOR dependency is linked to a lack of adaptive capability due to suppressed mRNA translation. **(a)** Distribution of PDOs along FMD trajectory (MOFA factor 2 loadings) highlighting extremes on the sensitivity axis that were chosen for perturbation experiments. **(b)** GSEA based on RNA-seq data showing normalized enrichment scores (NES) of pathways associated with mTOR dependency at baseline. Reactome pathways [33] were used for the pathway activity analysis. **(c)** Transcriptional and proteomic shift in Hallmark signatures [30], shown as the NES of mTOR-sensitive relative to resistant PDOs **(d)** Imputed activity of key regulatory kinases in sensitive compared to resistant PDOs. **(e)** Scatter plot showing the decoupling of PI3K-AKT-mTOR Hallmark signatures occurring between RNA and protein levels, expressed as log_2_ fold changes (LFC) between sensitive and resistant PDOs. **(f)** Baseline expression (variance-stabilizing counts, VST counts, left, normalized protein intensity right) of *ANXA1* and *PIK3AP1* in representative PDOs of both extremes of FMD axis. **(g)** NES of RNA-seq based GSEA of FMD, fetal and LGR5-ISC in PDOs following SN38 and mTOR inhibitor treatment. **(h)** Transcriptomic and proteomic response of sensitive and resistant PDOs to mTOR inhibition shown as NES after GSEA of Hallmark signatures and GO biological processes (relative to DMSO control). **(i)** Imputed kinase activity of key signaling and cell-cycle nodes following mTOR inhibition in both PDO groups, relative to DMSO controls. All p-values are calculated using Wilcoxon rank-sum test (two-sided).

Earlier characterization of the FMD signature established an axis of opposing stemness programs, where fetal-like signatures correlated with Factor 2 negative expression loadings (negative FMD score) and the homeostatic ISC-Lgr5 signature [21] was anti-correlated, peaking instead in the PI3K/mTOR independent Factor 2 positive (FMD score positive) loadings (Fig. 3c). Consistent with this depletion in adult stemness, phospho-proteomic profiling revealed increased GSK3B activity in PI3K/mTOR sensitive PDOs (Fig. 5d). Given its role as a negative regulator of the Wnt/β-catenin cascade [32], its high activity could provide an explanation for the loss of Lgr5+ driven stemness in these PDOs. This would suggest that mTOR dependency in CRC is an emergent feature of specific cells shifted into a fetal-like state. In this state, Wnt-driven homeostatic ISC signaling is suppressed and replaced by an mTOR-dependent developmental program.

To determine how the fetal-like state of mTOR-dependent PDOs changes upon drug perturbation with chemotherapy (SN38, active metabolite of pro-drug Irinotecan) and mTOR inhibition (Vistusertib), we quantified the enrichment of the FMD signature, fetal signatures and ISC-Lgr5 stemness signature across our representative cohort of PDOs at both transcriptomic and proteomic levels (Fig. 5g). Consistent with established literature regarding chemotherapy-induced selection of fetal-like plasticity in CRC [10], both mTOR-sensitive and-resistant PDOs demonstrated transcriptional enrichment of both FMD and fetal signatures in response to SN38 treatment. In contrast to the more generalized response seen with SN-38, mTOR inhibition using Vistusertib revealed a stark difference in the induction of stemness programs. While Vistusertib elicited a strong induction of both fetal signatures and FMD signature (with NES > 1) in the mTOR-sensitive models relative to their DMSO baseline, resistant models showed a repression of fetal signatures (NES < 0) under PI3K/mTOR pathway suppression.

Pathway analysis (GSEA) demonstrated that mTOR inhibition using Vistusertib engaged the intended target, indicated by the downregulation of mTORC1 signaling and ribosomal machinery (ribosome biogenesis and rRNA processing) on transcriptomic and proteome-level, respectively (Fig. 5h). This was mirrored at the functional level where MTOR kinase activity was suppressed in both sensitive and resistant PDOs (Fig. 5i). However, the two groups displayed diverging adaptive processes. Resistant PDOs induced compensatory mechanisms at RNA level, characterized by upregulation of translational machinery and DNA replication. This increased transcriptional activity correlated with increased activity of *CDK2* and suggests cell cycle induction. In contrast, mTOR-sensitive PDOs failed to meaningfully activate *CDK2*, instead exhibiting high *CHEK1* activity, a marker of replicative stress or an unresolved DNA damage response, ultimately leading to cell death.

### The PI3K/mTOR-dependent fetal CRC stem cell state is driven by an upregulated SRC signaling network

To further functionally dissect the signaling network driving the FMD phenotype, we conducted comprehensive profiling of 277 kinase inhibitors in the 8 PDOs of the validation cohort (Fig. 6a, Supp. Table 6). Kinase inhibitors were considered to have differential activity in the mTOR-sensitive PDO group if the median AUC difference between the groups exceeded 10% and reached a significant effect size based on the overall variance between the PDOs (Cohen’s d > 0.8). Out of the inhibitors profiled, mTOR-sensitive PDOs showed selective sensitivity to 14 compounds. Beyond mTOR inhibitors, these included inhibitors targeting Src family kinases (SFKs), specifically A419259 (Cohen’s d = 1.52, p = 0.057) and Saracatinib (Cohen’s d = 1.51, p = 0.057) and the selective BTK inhibitor Ibrutinib (Cohen’s d = 1.59, p = 0.057) (Fig. 6b).

**Fig 6.**
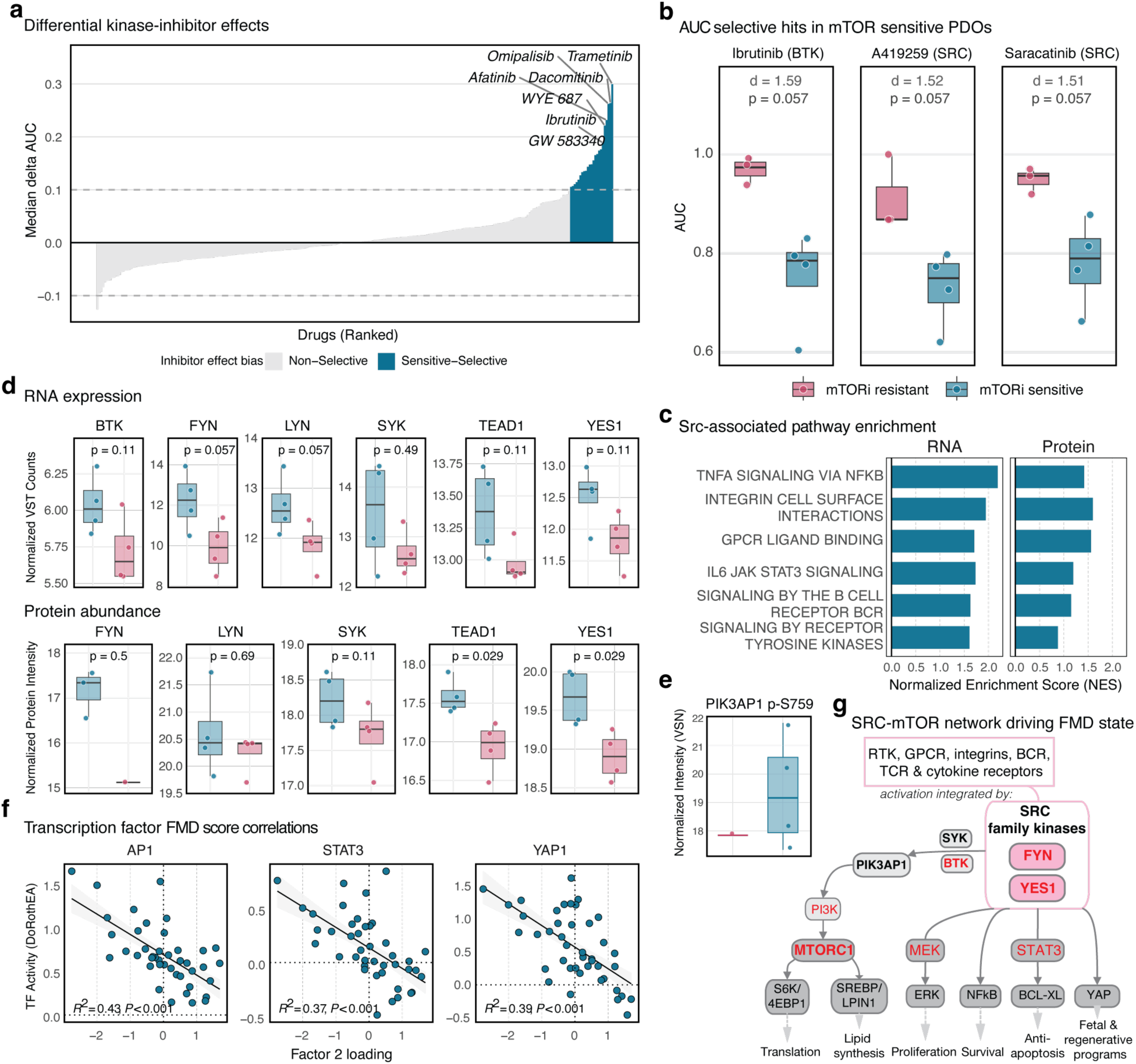
The mTOR dependent fetal CRC stem cell state is associated with an upregulated SRC signaling network. **(a)** Waterfall plot of kinase inhibitor profiling results, identifying compounds with selective efficacy in mTOR-sensitive PDOs (blue), ranked by the difference in median AUC between 4 PDOs from each extreme of the FMD axis. **(b)** Boxplots showing the AUC for inhibitors of SRC and BTK with selective activity in mTOR-sensitive PDOs. **(c)** GSEA based on RNA-seq data showing normalized enrichment scores (NES) of Hallmark and Reactome pathways associated with SRC pathways were used for the pathway activity analysis. **(d)** Baseline expression (VST counts) and protein abundance (normalized intensity) of SRC family kinases, associated kinases and downstream effector TEAD1. **(e)** Normalized intensity of PIK3AP1 phosphorylation at S759 in mTOR-sensitive and-resistant PDOs. **(f)** Correlation of DoRothEA transcription factor activity of SRC-and fetal-state-associated transcription factor with FMD score. **(g)** Proposed signaling network model illustrating SRC-mTOR axis in the FMD state. Model highlights how upstream effectors (growth factor and chemokine receptors, and integrins) converge on SRC family kinases (FYN, YES1) which engage PIK3AP1 as an amplifier to maintain PI3K-mTOR signaling to maintain translation alongside parallel proliferative and survival pathways (MAPK, STAT3, YAP). d denotes the effect sizes calculated using Cohen’s D. All p-values were calculated using Wilcoxon rank-sum test (two-sided).

To investigate how baseline differences between the two PDO groups were associated with this selective sensitivity, we performed GSEA on RNA and protein level. We identified several enriched pathways in both omics layers that have been linked to SRC activation, such as Integrin cell surface interactions, GPCR ligand binding and signaling by B-cell receptors and tyrosine kinases (Fig. 6c), for which SFKs act as signal transducers [34]. Further, additional pathways found downstream in the signaling cascade of SFKs were found to be upregulated, e.g. IL6-JAK-STAT signaling (Fig. 6c).

Molecular validation and investigation of leading-edge genes driving this enrichment of SFK-associated pathways confirmed the observed sensitivity was driven by specific SFK members, rather than c-Src itself. mTOR-sensitive PDOs displayed significantly higher baseline protein abundance of YES1 (p = 0.029) and TEAD1 (p = 0.029), along with elevated RNA expression of *LYN*, *FYN* and *BTK* (Fig. 6d). Activation of the SRC pathway in general has been linked to the regenerative fetal-like reprogramming involved in the tissue remodeling observed in CRC [19], but YES1 specifically has been reported to directly increase localization of YAP to the nucleus [35], where the presence of the transcription factor TEAD is required to allow DNA binding [36]. Beyond its effects on YAP, the activity of Src family kinases like YES1 appears to be required for MTORC1 activation [37]. As such, the co-expression of *TEAD1* and *YES1* may provide a mechanistic link to the fetal-like transcriptional programs enriched in the mTOR-sensitive PDOs.

To further explore the signaling network underlying this sensitivity, we examined the phosphorylation status of the key adapter protein PIK3AP1 enriched in the FMD signature, linking SFK signaling to PI3K activation (Fig 3b). Not only was PIK3AP1 protein more than 4-times more abundant in mTOR-sensitive PDOs (Fig. 5f), but we also detected phosphorylation of PIK3AP1 at Serine 759 (Fig. 6e), which is a PIK3AP1 modification that has been detected upon B-and T-cell receptor activation, followed by upregulation of the PI3K/mTOR signaling pathway [38]. During B-cell receptor signaling - one of the pathways found enriched in mTOR-sensitive PDOs both on protein and RNA level (Fig. 6c)-phosphorylation of PIK3AP1 by BTK or SYK is required to provide a binding site for the p85 subunit of PI3K, thereby promoting PI3K activation [31,39]. Interestingly, both *BTK* and *SYK* expression were elevated in mTOR-sensitive PDOs (Fig 6d), and BTK inhibition selectively affected mTOR-sensitive PDOs, suggesting a pivotal role for BTK-mediated processes in the survival of mTOR-dependent PDOs.

Recent studies report a much broader role for PIK3AP1, beyond its ability to relay B-Cell receptor signaling to PI3K, in primary immune response, where it has been linked to toll-like receptors (TLRs) as well as interleukin-1 (IL-1) family receptors [40]. Considering that both interferon and interleukin signaling were found to be upregulated in mTOR-sensitive PDOs (Fig. 5b), our data indicates that PIK3AP1 could serve as a central signaling integration node in the SFK-mTOR survival axis.

Finally, we assessed the activity of transcription factors downstream of this hub. We found that the activity of AP1, STAT3 and YAP1 strongly correlated with the FMD score (R^2^ of 0.43, 0.37 and 0.39 respectively, Fig. 6f) in the original discovery cohort of 42 PDOs. Its negative end represents the fetal-like mTOR dependency state. Notably, these transcription factors have been not just been associated with SRC signaling, but more recently also with the plastic state fetal-like stem cells in CRC [10,14]. This suggests that the YES1-YAP1 and SFK-AP-1 axes are the primary drivers of the regenerative, fetal-like stemness program in the fetal mTOR-dependency observed in these PDOs. Altogether, these findings support a model in which Src-YAP and BCR-like signaling (via BTK and SYK) converge on PIK3AP1 in fetal-like mTOR-dependent PDOs. This network may sustain PI3K/mTOR signaling and fetal-like programs despite the reduced mTOR-mediated translational activity (Fig. 6g).

## Discussion

Our study provides a comprehensive multi-omics and functional analysis of colorectal cancer stem cell states by integrating transcriptomic, genomic and large-scale drug perturbation profiling of a large PDO cohort and extensively validating these findings in patients’ tumors and organoid datasets. Thereby, we define the landscape of CRC adult and fetal/regenerative stem cell states and identify potentially tractable dependencies, such as an SRC/mTOR axis that we further explored by proteomics and functional kinase inhibitor profiling.

Recent single-cell atlases have substantially advanced the understanding of cellular heterogeneity and developmental plasticity in CRC (see review [5]) and have identified different stem cell states in CRCs, such as proliferative and revival cancer stem cells [9] or organoid developmental trajectories organized along MAPK activity gradients [11,12]. Yet, these approaches largely remain descriptive and provide limited insight into therapeutically actionable dependencies and mechanisms behind this. Hence, linking cell states to functional perturbations is an important task [13]. Using a large cohort of patient-derived organoids combined with systematic drug screening, we systematically identified continuous developmental trajectories ranging from adult intestinal stem cell-like to regenerative/fetal-like states that are conserved across organoid, bulk tumor, and single-cell datasets (Fig. 4). We could show that fetal/regenerative features are inherent cellular states as complimentary to previous studies showing such states being activated upon external stimuli [14]. These findings support the concept that CRC stem cell states are not discrete entities but rather exist along dynamic developmental continua shaped by cellular plasticity and regeneration/fetal-plasticity-associated programs.

Importantly, our data demonstrate that functional phenotyping can uncover biologically and therapeutically relevant vulnerabilities that are not readily identifiable by transcriptomic profiling alone. Although the FMD trajectory displayed elevated mTOR-associated transcriptional programs, proteomic analyses revealed suppression of translation-associated processes (Fig. 5), suggesting a disconnect between transcriptional pathway activation and protein translation. This is consistent with the well-established observation that mRNA abundance does not necessarily predict protein abundance, which is also dependent on translation levels, protein stability and turnover [41]. In particular, pathways related to RNA processing and translation have previously been reported to show comparatively low RNA-protein concordance [42].

In this context, the pronounced sensitivity of FMD-high PDOs to PI3K/mTOR inhibition, particularly to the dual mTORC1/2 inhibitor Vistusertib and the PI3K/mTOR inhibitor Gedatolisib suggests that these PDOS depend on a constrained and therapeutically vulnerable mTOR signaling state. One possible interpretation is that the transcriptional up-regulation of mTOR-associated programs reflects a compensatory response to reduced translational pathway output at the proteome level. Accordingly, further pharmacological suppression of this axis may exceed the adaptive capacity of fetal-like CRC cells, leading to selective sensitivity to PI3K/mTOR pathway inhibition.

Beyond a PI3K/mTOR dependence, we further linked the FMD score to an activated SRC-associated signaling network acting upstream of mTOR signaling. The co-sensitivity of mTOR-dependent PDOs to SRC family kinase and BTK inhibitors suggests that upstream SRC/BTK-associated signaling contributes to the maintenance of the FMD state. Consistent with this, our integrated drug-profiling, transcriptomic, proteomic and phosphoproteomic analyses support a model in which SRC/YAP-associated [43] and BCR-like signaling converge on PIK3AP1, thereby sustaining PI3K/mTOR pathway activity [31,39] and fetal-like transcriptional programs despite reduced translation-associated protein output. This is notable in the light of the renewed interest in SRC family kinases, YES1 in particular, as mediators of therapy resistance [43,44]. Although SRC inhibitors have historically shown limited efficacy as monotherapies in solid tumors, our data suggest that fetal-like, mTOR-dependent CRC models may represent a more defined molecular context in which SRC/BTK-associated signaling contributes to a targetable adaptive state.

Together, these findings suggest that fetal-like mTOR-dependent CRC PDOs are not broadly therapy-resistant but instead occupy a specialized signaling state with distinct targetable dependencies. Disruption of either upstream SRC/BTK-associated inputs or downstream PI3K/mTOR signaling may therefore compromise the adaptive signaling network that supports this state. More broadly, these data highlight how integrated functional perturbation approaches can refine molecular CRC classifications by resolving mechanistically distinct and therapeutically actionable subgroups within tumor stem cell states.

Clinical targeting of the PI3K/mTOR pathway in colorectal cancer has so far shown limited efficacy, despite frequent activation of this signaling axis in CRC [45]. This may partly reflect the biological heterogeneity and plasticity of CRC, as well as the absence of biomarkers enabling patient stratification. Our data suggest that sensitivity to PI3K/mTOR inhibition may not be broadly linked to canonical pathway activation by mutations but rather associated with a distinct regenerative/fetal-like tumor cell state characterized by impaired translational adaptability and SRC-associated signaling. These findings may provide a potential explanation for the limited success of PI3K/mTOR-directed therapies in CRC and support the concept that developmental tumor states could inform future stratification approaches for combination therapy.

Some limitations of our study should be acknowledged. First, although patient-derived organoids preserve key epithelial tumor-intrinsic programs, they incompletely capture interactions with the tumor microenvironment, which may additionally shape stem cell plasticity and therapeutic response. In this context, interpretation of bulk tumor transcriptomes remains challenging due to admixture with stromal and immune cell populations, as previously noted in colorectal cancer classification studies. Our integration of single-cell datasets partially addresses this limitation by confirming that the identified developmental gradients are present within malignant epithelial compartments. Additionally, while our analyses integrate large-scale organoid perturbation profiling with validation across multiple bulk and single-cell CRC cohorts, the identified SRC/mTOR dependencies were not validated *in vivo* and therefore require further investigation in physiologically more complex model systems.

In conclusion, our study establishes a functional and molecular landscape of continuous CRC stem cell states by integrating patient-derived organoid drug perturbation profiling with transcriptomic analysis, as well as single-cell data from CRCs. Beyond descriptive cellular atlases, this approach identifies mechanistic backgrounds of cell state trajectories. In particular, we define a regenerative/fetal-like CRC state characterized by SRC-associated PI3K/mTOR dependency and impaired translational adaptability. These findings demonstrate how functional profiling can complement molecular classification approaches to uncover biologically relevant and potentially targetable tumor states in CRC.

## Methods

### Patients

All patients were recruited at University Hospital Mannheim, Heidelberg University, Mannheim, Germany. We included patients diagnosed with microsatellite stable (MSS) colon and rectum cancer in this study and obtained biopsies from their primary tumors via endoscopy or from metastatic lesions by ultrasound-guided fine-needle biopsy. Exclusion criteria were active HIV, HBV or HCV infections. Clinical data, tumor characteristics and molecular tumor data were pseudonymized and collected in a database. The clinical data of the participants of this paper can be found in Supp. Tab. 1. The research was approved by the Medical Ethics Committee II of the Medical Faculty Mannheim, Heidelberg University (Reference no. 2014-633N-MA and 2016-607N-MA). All patients gave written informed consent before tumor biopsy was performed.

### Organoid culture

Organoids were established from tumor biopsies as previously reported [46]. Briefly, biopsies were washed and digested with Liberase TH (Roche) before embedding into Matrigel (Corning) or BME (Cultrex). Advanced DMEM/F12 (Thermo Fisher Scientific) medium with Pen/Strep (Gibco), Glutamax (Gibco) and HEPES (Gibco) was supplemented with 100 ng/ml Noggin (PeproTech), 1 x B27 (Thermo Fisher Scientific), 1,25 mM n-Acetyl Cysteine (Sigma), 10 mM Nicotinamide (Sigma), 50 ng/ml human EGF (Peprotech), 10 nM Gastrin (Peprotech), 500 nM A83-01 (Biocat), 10 nM Prostaglandin E2 (Santa Cruz Biotechnology), and 100 mg/ml Primocin (Invivogen). 10 µM Y-27632 (Selleck chemicals) were added after thawing and passaging. Organoids were passaged every 7–10 days and medium was refreshed every 2–3 days.

### Drug profiling

Drug profiling was performed in similar way as previously published with few adaptations [46,47]. For cell seeding, organoids were incubated with TrypLE Express (Gibco) at 37°C until small clusters and single cells were obtained. Chemical separation was supported by mechanical shearing using a 1000 µl pipette and digestion was visually controlled by light microscopy. Organoid fragments were filtered through a 40 µm strainer (pluriSelect) to prevent large organoid clusters and afterwards quantified. Culture medium was supplemented with Y-27632 and growth factor-reduced BME type 2 was added to a concentration of 0.75 mg/ml (seeding medium). For seeding, organoids were resuspended in 50µl of seeding medium and seeded into 384 well plates with a multidrop dispenser (Thermo Fisher Scientific). Drug treatment was performed on day 3 after seeding. Medium was aspirated, discarded and 45 µl fresh medium was added, drugs were pre-diluted in medium and 5 µl of diluted drugs were added. All pipetting steps were performed by a Biomek FX^P^ robotic device (Beckman Coulter). Viability was measured on day 7 after seeding. Medium was aspirated and discarded before 30 µl undiluted CellTiter-Glo^®^ solution (Promega) was added to each well. After 30 minutes of incubation at room temperature, luminescence was measured by a Mithras reader (Berthold Technologies).

### Drug libraries

Two clinical libraries were used for drug profiling in two sub-cohorts: A 140-drug clinical library for the validation cohort of 22 individual PDOs [47] contains a comprehensive selection of FDA-approved cancer-targeting small molecule drugs (excluding antibodies, antibody-drug-conjugates as well as drugs mainly targeting the immune system or tumor microenvironment). A second cohort for discovery of 56 individual PDOs, 62-drugs clinical library [46] contains a focused subset of this collection. Drugs within the clinical libraries and their maximum concentrations were selected individually based on literature review of 2D and 3D cell culture assays, as well as own previous experiments. Additionally, we used a kinase-inhibitor library with 277 compounds to perform perturbation-based mapping of signaling pathways. This library is depicted in Supplementary Table 6. In all libraries, each compound was screened in five different concentrations after each five-fold dilutions. 5 µM bortezomib was used as positive control, DMSO as negative control. All libraries were arranged in a random layout using a Biomek NX^P^ robotic system (Beckman Coulter). Drugs were purchased from Selleck Chemicals or MedChemExpress.

### DNA sequencing

Mutations were analyzed primarily by whole exome sequencing using the DKFZ-OTP pipeline [48] with a subset of data from rectal cancer organoids previously published [47]. Another set of previously published lines had already been characterized by targeted amplicon sequencing as described earlier [46]. DNA was isolated using the DNeasy Blood and Tissue kit (Qiagen). Variants were annotated with ANNOVAR [49] and only exonic or splicing mutations classified as “frameshift deletion”, “frameshift insertion”, “nonsynonymous SNV”, “stopgain” or “stoploss”, with an allele frequency >0.1, which were present in COSMIC database, were considered for further analysis.

### Proteomic and phospho-proteomic profiling

Proteomic analyses of colorectal cancer patient-derived organoids (PDOs) were performed essentially as previously described [47]. Briefly, organoid pellets were lysed in guanidine hydrochloride-containing lysis buffer, sonicated, and subjected to overnight tryptic digestion. Peptides were purified using the SP3 clean-up protocol and analyzed by nanoLC–MS/MS using an EASY-nLC 1200 system coupled to a Q Exactive HF Orbitrap mass spectrometer (Thermo Fisher Scientific) operated in DIA mode. Raw files were processed using Spectronaut (Biognosys) applying a directDIA workflow against the UniProt human proteome database. Protein quantification and normalization were performed using the integrated Spectronaut analysis pipeline. To account for the inherent heteroscedastic nature of proteomic data, all intensities were adjusted using variance stabilizing normalization (VSN) available in the R package vsn (v3.76.0).

### Expression profiling

Organoid RNA was isolated with the RNeasy kit (Qiagen) following the manufacturer’s instructions. Organoids were pelleted by centrifugation and frozen in RLT buffer containing 1% β-mercaptoethanol before RNA isolation.

RNA-seq was performed with the TruSeq stranded protocol (Illumina) by the DKFZ Sequencing Core Facility and data were processed using the DKFZ OTP pipeline [48]. For microarray analysis, samples were hybridized on Affymetrix Human Genome U133 plus 2.0 arrays (Affymetrix).

### Transcriptomics data processing

For in-house PDO biobank, raw CEL files were processed using affy (v1.86.0) with Robust Multi-array Average (RMA) normalization (background correction, quantile normalization, and probe summarization). Based on initial principal component analysis (PCA) of in-house PDO microarray data, samples were moderately separated by their gender (Supp. Fig. 1b). Similarly, X-or Y-linked genes were moderately correlated with PC1 axis (Pearson r(PC1, X) =-0.4 or r(PC1, Y) = 0.3). We adjusted for gender to isolate epithelial biology and modeled expression values with gender as covariate using removeBatchEffect() function in limma (v3.64.3). We utilized batch-corrected expression values for the downstream analysis.

Raw microarray data from external PDO cohorts (Gene Expression Omnibus; GSE64392, GSE74843, GSE128213) were processed from CEL files using RMA normalization implemented in oligo (v1.72.0) and affy (v1.86.0). Samples were restricted to CRC organoids derived from primary tumors and MSS cases. To enable integration across datasets, gene expression matrices were combined and batch effects due to cohorts were corrected using the ComBat algorithm implemented in sva (v3.56.0). The resulting normalized and batch-corrected expression matrix was used for downstream analyses.

Gene expression data for TCGA-COAD and TCGA-READ cohorts were obtained from The Cancer Genome Atlas (TCGA) ([50], https://www.cancer.gov/tcga). Analyses were restricted to primary tumors classified as MSS based on MANTIS scores (MSS defined as ≤ 0.4 as suggested by Kautto et al. [51]). Expression values were transformed as log₂(TPM + 1). Consensus Molecular Subtype (CMS) annotations [2] were obtained from published classifications and matched to TCGA samples. KRAS mutation status was derived from TCGA somatic mutation data and integrated for downstream association analyses.

For single-cell analyses, we used the CRC atlas from Chu et al. [52], restricting the data to datasets (n = 6 from four distinct studies, [26–29]) with sufficient malignant cell representation (>1,000 malignant cells per dataset) and excluding polyp samples. As preprocessing and quality control had been performed by the original authors, no additional QC filtering was applied, except for the removal of genes expressed in fewer than two cells. For analyses involving projection onto epithelial axes or marker expression, expression values were z-scaled in Seurat (v5.3.1) with regression of total UMI counts to account for differences in library sizes.

### Diffusion maps

We computed diffusion maps using the DiffusionMap() function from destiny (v3.22.0) with default parameters on normalized gene expression values. For the in-house PDO cohort, we used RMA-normalized and batch-corrected expression values. For external PDO datasets, we used integrated RMA-normalized expression values. For TCGA data, we used log_2_(TPM+1)-transformed expression values.

The gene selection was based on PC loadings derived from prior dimensionality reduction in the in-house cohort. Specifically, we selected mutually exclusive top 50 and bottom 50 genes ranked by loadings for each of PC1 and PC2 axes, capturing genes with the strongest positive and negative contributions to each axis.

Diffusion pseudotime was inferred using the DPT() function, specifying a single root tip (tips = 1). The first diffusion pseudotime component was extracted and used as the sample-level pseudotime measure for downstream analyses.

### Permutation-based assessment of trajectory robustness

To assess whether the observed trajectories could arise by chance, we generated diffusion maps from 100 random genes matched in size to the PC-derived gene sets 1000 times. For each random gene set, we calculated the Spearman correlation between pseudotime and the first diffusion component, DC1, generating a null distribution of pseudotime–DC1 correlations (Supp. Fig. 1d). The observed correlations for the PC1-and PC2-derived epithelial gene sets were then compared against this null distribution.

### Evaluation of gene expression modules

To contextualize our epithelial biology and drug sensitivity axes with respect to prior studies, we applied multiple published gene expression signatures from diverse biological contexts: i) fetal/regenerative state signatures (Supp. Tab. 3), ii) Hallmark gene sets [30], and iii) CRC-specific gene signatures (Supp. Tab. 4).

To further characterize the biological programs associated with the extremes of each axis, we performed functional enrichment analysis on genes with the strongest loadings. We selected the top 200 genes with the highest positive loadings and the top 200 genes with the most negative loadings from the PC1 or PC2 loadings. Enrichment analysis was then performed separately for the positive and negative gene sets using gost() from gprofiler2 (v0.2.3), testing for over-representation in GO Biological Process [53], KEGG [54], and Reactome [33] pathways. Multiple testing correction was performed using FDR, and terms with adjusted *p* < 0.05 were considered significant.

To represent the gene expression modules (e.g., the FMD score, PC1 and PC2 axes, and published gene signatures), we computed module scores by taking the mean z-scaled expression values (mean = 0, standard deviation SD = 1 across samples for each gene) of all genes within each gene set for each sample.

### Multi-omics analysis

To integrate gene expression, drug response, and mutation data across the first CRC PDO cohort (n = 42), we used multi-omics factor analysis (MOFA2 v1.13.0, [15]). For the expression module, we selected the top 10% most variable probes based on the coefficient of variation and used the average log2-transformed expression calculated per PDO and gene. For the drug response module, we averaged DMSO-normalized AUC values per PDO–drug pair. For the mutation module, we merged hard-filtered exome (allele frequency of > 10%) and amplicon mutation calls, retained genes mutated in at least three samples and encoded mutation values as binary variables.

These three modalities were combined into a single MOFA model with Gaussian likelihoods for expression and drug response data and a Bernoulli likelihood for mutation data. The model was initialized with 7 latent factors and trained using default settings with slow convergence mode, stochastic inference disabled, and a fixed random seed of 42. Factors explaining less than 1% of variance per view were removed during training.

### Survival analysis

Progression-free survival (PFS) and overall survival data (OS) data were obtained from TCGA COAD/READ PanCancer Atlas clinical annotation. Survival times were measured in months. We excluded samples with missing survival status from the analyses.

Kaplan-Meier survival curves were generated using the survfit() function from survminer (v0.5.1), including 95% confidence intervals (CI) and risk tables where appropriate.

Univariable Cox proportional hazards regression was performed for PFS using the coxph() function from survival (v3.8-3). Each variable was tested independently, including F2 score, age, sex, and pathological stage. Hazard ratios (HRs), 95% confidence intervals (CIs), and Wald test P values were reported for each variable individually.

Multivariable Cox proportional hazards regression was used to evaluate whether F2 score was independently associated with PFS after adjustment for clinical covariates. The model included F2 score, age, sex, and pathological stage as covariates. Results are reported as adjusted HRs with 95% CIs and p values. HRs were displayed on a logarithmic scale. P values below 0.05 were considered statistically significant.

## Statistical analyses

Associations between molecular features and diffusion components (DC1 and DC2) were assessed using linear regression models. Standard linear regression models were used to quantify the relationship between sample annotations (e.g., tumor site, histological subtype, mutational status) and diffusion components (DC1 and DC2). Model fit was assessed using the coefficient of determination (R²), and statistical significance of individual predictors was evaluated using two-sided tests.

### Gene set enrichment analysis

Gene set enrichment analysis was performed across multiple datasets. For specialized custom gene sets, including fetal and Lgr5+ signatures, fast pre-ranked GSEA was executed using the fgsea R package (v1.34.2). Standardized functional annotations were evaluated using the clusterProfiler ecosystem (v4.16.0), specifically ReactomePA (v1.52.0) was used to analyze Reactome pathways and Gene Ontology (GO) Biological Processes (BP) were assessed via gseGO (mapped using org.Hs.eg.db v3.21.0) and MSigDB Hallmark Pathway collections [30] were accessed using the universal GSEA() function (MSigDBr, v25.1.1). For all implementations, inputs were either ranked by factor loadings or by differential expression metrics (Waldman Statistic from DESeq2 (v1.48.2), or t-statistic from limma (v3.64.3)) and a significance threshold of p < 0.05 was applied. Statistical significance was assessed by applying the Benjamini–Hochberg procedure to control the false discovery rate (FDR). Hallmark and GO enrichment analysis were restricted to gene sets containing between 10 and 500 genes.

### Imputation of Kinase Activity Scores

Raw phospho-proteomic intensities were filtered to retain phosphosites with less than 20% missing values across samples, followed by variance-stabilizing normalization (VSN). Duplicate gene-phosphosite combinations were summed prior to downstream analysis. Differential phosphorylation analysis was performed using the limma package (v) in R. A linear model incorporating treatment condition and PDO as covariates was fitted to account for inter-organoid variability. Phosphosites were required to have at least three non-missing values in each condition being compared. Empirical Bayes moderation with trend and robust fitting was applied, and t-statistics from the resulting contrasts were used as input for downstream activity inference. Kinase activity was estimated using the weighted mean enrichment method (1000 permutations) implemented in the decoupleR package (v2.14.0) with a kinase-substrate network derived from OmniPath. Only phosphorylation and dephosphorylation interactions were retained and any ambiguous kinase-substrate pairs with conflicting modes of regulation were removed. Kinases with fewer than five measured sites were excluded from activity scoring. Pathway-level activity was then inferred using the PTMSigDB database (v2.0.0, accession: 12.07.2024). Phosphosite identifiers were mapped from gene symbols to UniProt accessions using the org.Hs.eg.db package (v3.21.0) and pathway activities were estimated using the same weighted mean enrichment approach. Pathway significance was assessed by permutation-derived p-values and a composite score integrating activity magnitude and statistical significance (|score ×-log10(p)|) was used to prioritize results.

### Immunofluorescence staining and imaging

Embedded PDOs were fixed in 4% Paraformaldehyde, blocked with 1% BSA and incubated with primary antibodies for ANXA1 (ab214486, Abcam) and OLFM4 (#MAB9218, R&D Systems) [55], followed by incubation with appropriate secondary antibodies and nuclear counterstaining with Hoechst 33324. Images were acquired using an Incell Analyzer 6000 microcope (GE Healthcare) at 20x magnification and processed identically across compared conditions using FIJI (v 2.17.0/1.54p).

## Data availability

The datasets generated in this study are not yet publicly available at the time of preprint posting. Expression profiling data will be deposited in the Gene Expression Omnibus (GEO). Proteomics data will be deposited in the PRIDE repository and made available through ProteomeXchange. Human next-generation sequencing data from organoid models will be deposited or made available through controlled-access repositories, including the European Genome-phenome Archive (EGA) and/or the German Human Genome-Phenome Archive (GHGA), where applicable. Accession numbers will be provided in the final published version of the manuscript. Access to controlled human sequence data will be subject to review and approval by the responsible data access committees at the University Medical Center Mannheim and the German Cancer Research Center (DKFZ), completion of an appropriate data transfer or data use agreement, and compliance with the EU General Data Protection Regulation. Published datasets included in this study are available from the original sources cited in the manuscript.

## Contributions

ThM and BA contributed to conceptualization, methodology, formal analysis, writing of the original draft, and review and editing of the manuscript. ZL, LS, NR, LT, PS, EK, PA, JER, YP, TiM, EB, TZ, LD, NS and SB contributed to investigation. SB, ME, and JB contributed resources. SW contributed to supervision. MB and MPE contributed to supervision and funding acquisition. JB contributed to conceptualization, supervision, funding acquisition, resources, writing of the original draft, and review and editing of the manuscript. All authors reviewed and approved the final manuscript.

## Acknowledgements

We thank the NGS Core Facility of the German Cancer Research Center for help with exome sequencing and the Omics-IT Facility of DKFZ for help with sequencing data analysis. We thank the DKFZ Microarray core facility for performing expression profiling experiments and the Proteomics Core Facility of the DKFZ for performing Mass-Spectrometry-based proteomics profiling. We thank Shrihar Kanikar, Frank Herweck, Gabriela Atanasova and Kauthar Srour for excellent technical support with organoid establishment and biobanking. We thank Maryna Udovychenko, Ayse Civanna and Bianca Huber for excellent support in patient recruitment and the staff of the Central Endoscopy Unit Mannheim for assistance with tumor biopsies. This project was supported by the Hector Foundation II, the German Cancer Consortium DKTK Joint Funding Project Organoid Platform, the German Research Foundation (DFG) Grant FOR5806 (project number 537604907), the DKFZ-MOST program, as well as the Health + Life Science Alliance Heidelberg Mannheim that received state funds approved by the State Parliament of Baden-Württemberg. We further acknowledge support from the DKFZ International PhD Program (P.A.), the Oversea study program of the Guangzhou Elite Project (Z.L.), and the German National Academic Foundation (J.E.R.).

