## Supplementary material for "Multi-omics, organoid-based modeling reveals an SRC/mTOR-dependent fetal-like stem cell trajectory in colorectal cancer": Supp. Fig.

### Supplementary Figures

A

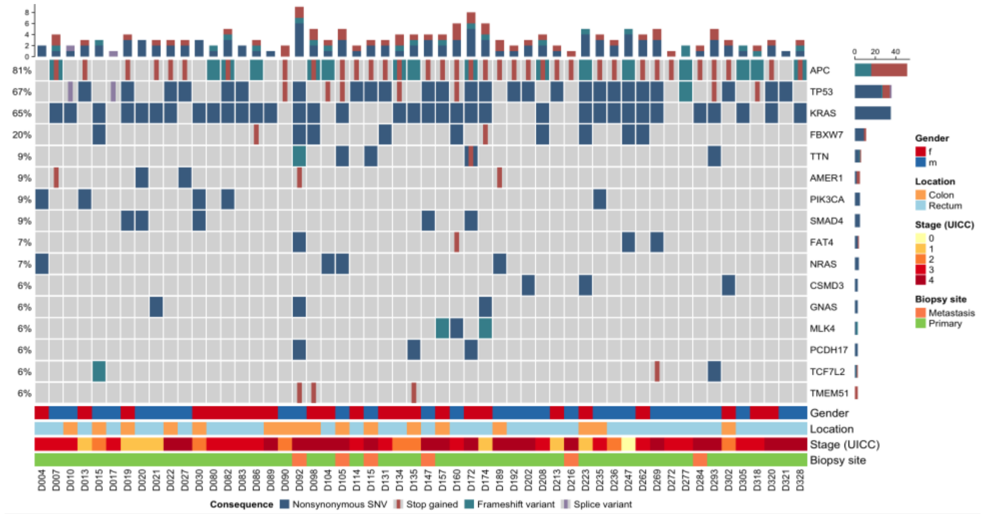


b

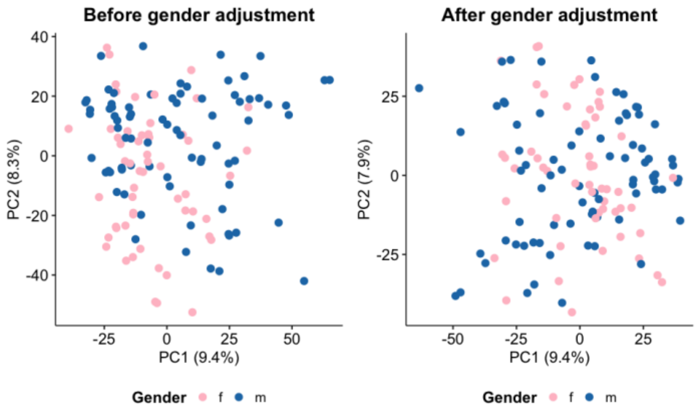


c

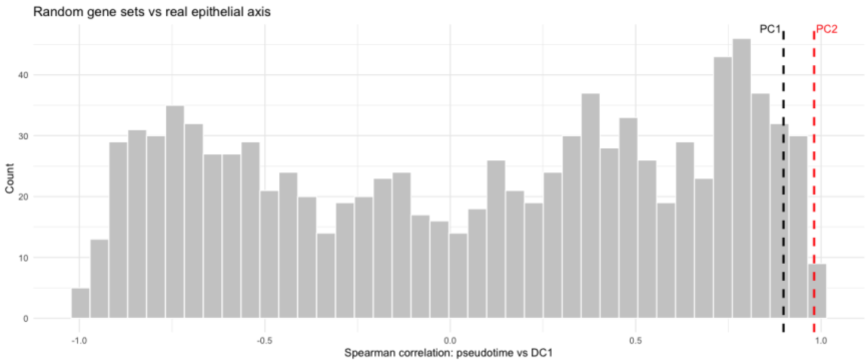

**Supp. Fig. 1. Cohort characterization, correction of confounding effects, and validation of epithelial plasticity axes. (a)** Oncoprint summarizing the mutational landscape of the PDO cohort, including recurrent alterations. Annotations indicate mutation types, patient characteristics (e.g., gender), and sample origin. **(b)** Principal component analysis (PCA) of PDO samples before (left) and after (right) adjustment for gender. Removal of gender-associated variation reduces clustering driven by sex and preserves biologically relevant structure. **(c)** Null distribution of Spearman correlation coefficients between diffusion component 1 (DC1) and pseudotime, generated from random gene sets (each 100 genes). The observed correlations for epithelial plasticity axes (PC1 and PC2; dashed lines) lie outside the null distribution, indicating that the epithelial gene programs capture non-random, biologically meaningful structure.

**a**

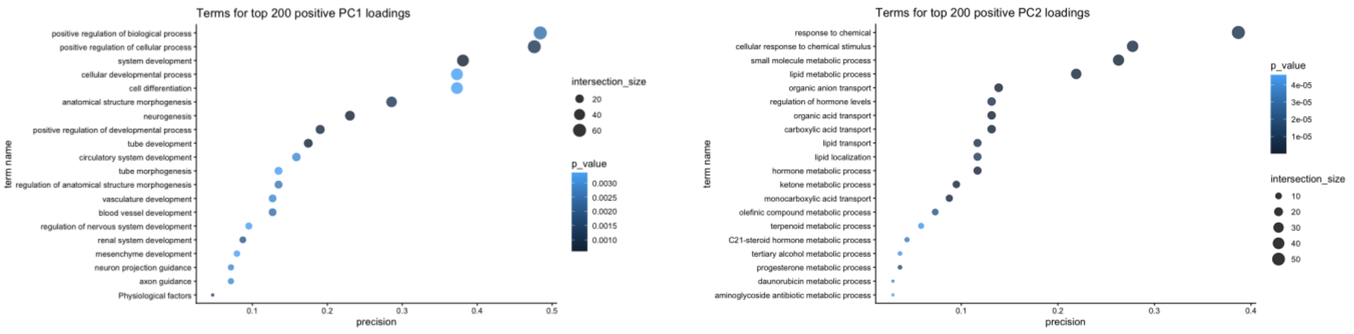
Terms for top 200 positive PC1 loadings
positive regulation of biological process
positive regulation of cellular process
system development
cellular developmental process
cell differentiation
anatomical structure morphogenesis
neurogenesis
positive regulation of developmental process
tube development
circulatory system development
tube morphogenesis
regulation of anatomical structure morphogenesis
vasculature development
blood vessel development
regulation of nervous system development
renal system development
mesenchyme development
neuron projection guidance
axon guidance
Physiological factors
term name
precision
0.1
0.2
0.3
0.4
0.5
intersection_size
20
40
60
p_value
0.0030
0.0025
0.0020
0.0015
0.0010
Terms for top 200 positive PC2 loadings
response to chemical
cellular response to chemical stimulus
small molecule metabolic process
lipid metabolic process
organic anion transport
regulation of hormone levels
organic acid transport
carboxylic acid transport
lipid transport
lipid localization
hormone metabolic process
ketone metabolic process
monocarboxylic acid transport
olefinic compound metabolic process
terpenoid metabolic process
C21-steroid hormone metabolic process
tertiary alcohol metabolic process
progesterone metabolic process
daunorubicin metabolic process
aminoglycoside antibiotic metabolic process
term name
precision
0.1
0.2
0.3
0.4
p_value
4e-05
3e-05
2e-05
1e-05
intersection_size
10
20
30
40
50

**b**

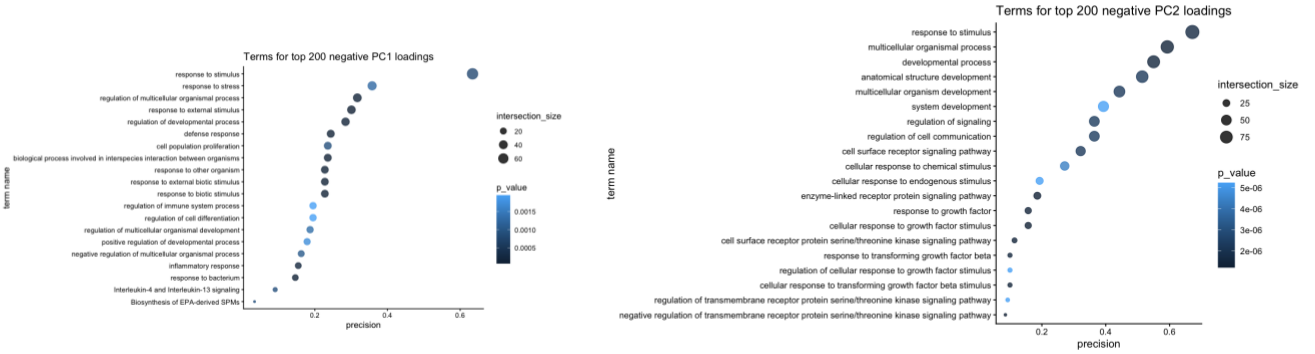
Terms for top 200 negative PC1 loadings
response to stimulus
response to stress
regulation of multicellular organismal process
response to external stimulus
regulation of developmental process
defense response
cell population proliferation
biological process involved in interspecies interaction between organisms
response to other organism
response to external biotic stimulus
response to biotic stimulus
regulation of immune system process
regulation of cell differentiation
regulation of multicellular organismal development
positive regulation of developmental process
negative regulation of multicellular organismal process
inflammatory response
response to bacterium
Interleukin-4 and Interleukin-13 signaling
Biosynthesis of EPA-derived SPMs
term name
precision
0.2
0.4
0.6
intersection_size
20
40
60
p_value
0.0015
0.0010
0.0005
Terms for top 200 negative PC2 loadings
response to stimulus
multicellular organismal process
developmental process
anatomical structure development
multicellular organism development
system development
regulation of signaling
regulation of cell communication
cell surface receptor signaling pathway
cellular response to chemical stimulus
cellular response to endogenous stimulus
enzyme-linked receptor protein signaling pathway
response to growth factor
cellular response to growth factor stimulus
cell surface receptor protein serine/threonine kinase signaling pathway
response to transforming growth factor beta
regulation of cellular response to growth factor stimulus
cellular response to transforming growth factor beta stimulus
regulation of transmembrane receptor protein serine/threonine kinase signaling pathway
negative regulation of transmembrane receptor protein serine/threonine kinase signaling pathway
term name
precision
0.2
0.4
0.6
intersection_size
25
50
75
p_value
5e-06
4e-06
3e-06
2e-06

**c**

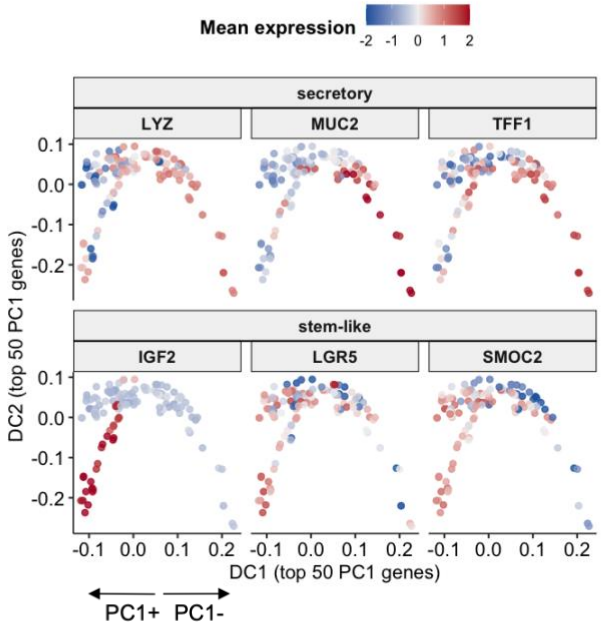
Mean expression
-2
-1
0
1
2
secretory
LYZ
MUC2
TFF1
stem-like
IGF2
LGR5
SMOC2
DC2 (top 50 PC1 genes)
0.1
0.0
-0.1
-0.2
-0.1
0.0
0.1
0.2
DC1 (top 50 PC1 genes)
PC1+
PC1-

**d**

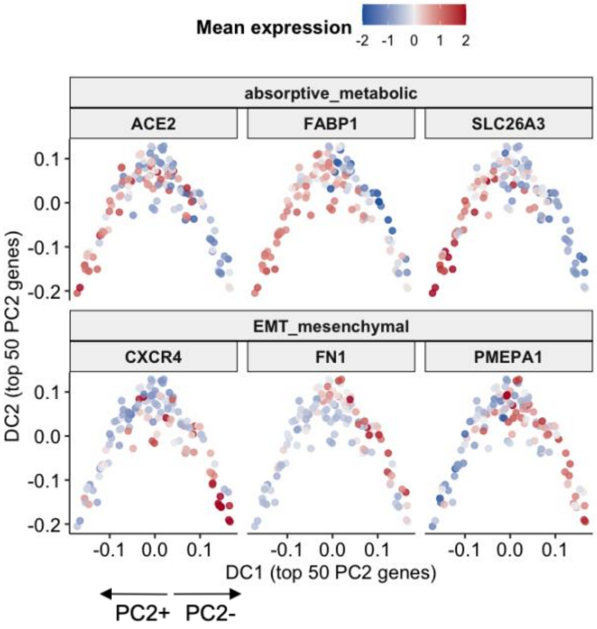
Mean expression
-2
-1
0
1
2
absorptive_metabolic
ACE2
FABP1
SLC26A3
EMT_mesenchymal
CXCR4
FN1
PMEPA1
DC2 (top 50 PC2 genes)
0.1
0.0
-0.1
-0.2
-0.1
0.0
0.1
DC1 (top 50 PC2 genes)
PC2+
PC2-

**Supp. Fig. 2. Functional annotation and biological interpretation of epithelial plasticity axes. (a)** Gene Ontology (GO) enrichment analysis of the top 200 positively (left) and negatively (right) loaded genes along PC1. Positively loaded genes are enriched for developmental and differentiation-related processes, whereas negatively loaded genes are associated with stimulus response, immune regulation, and stress-related pathways. Dot size represents gene set overlap (intersection size), and color indicates statistical significance (p-value). **(b)** GO enrichment analysis of the top 200 positively and negatively loaded genes along PC2. Positive PC2 loadings are enriched for metabolic and transport-related processes, including lipid and organic acid metabolism, while negative PC2 loadings are associated with signaling, cellular communication, and response to stimuli. **(c)** Expression of representative marker genes projected onto the diffusion map constructed using the top 50 PC1 genes. Z-scaled expression values for marker genes are shown from low (blue) to high (red). Distinct epithelial programs are captured along the axis, including secretory (LYZ, MUC2, TFF1) and stem-like (IGF2, LGR5, SMOC2) states, demonstrating the biological relevance of PC1. **(d)** Expression of representative marker genes projected onto the diffusion map constructed using the top 50 PC2 genes. Z-scaled expression values for marker genes are shown from low (blue) to high (red). Marker genes highlight absorptive/metabolic (ACE2, FABP1, SLC26A3) and EMT/mesenchymal (CXCR4, FN1, PMEPA1) programs, confirming that PC2 captures orthogonal epithelial phenotypes.

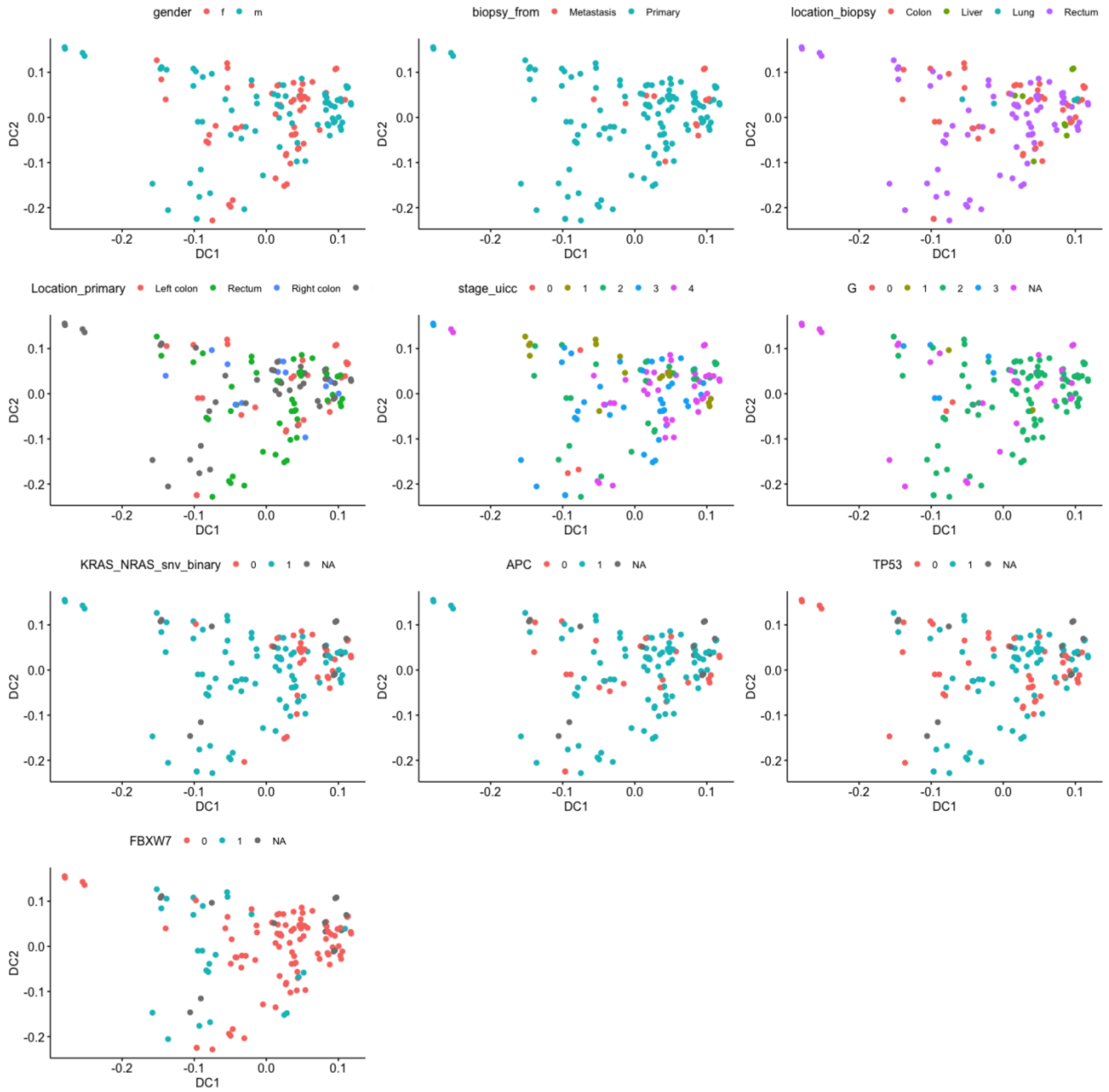


**Supp. Fig. 3. Clinical and genomic features on the epithelial plasticity axes.** Diffusion map based on top 50 highest and lowest PC1 and PC2 probes of PDO samples (DC1 vs DC2) colored by clinical variables (gender, biopsy origin and location, stage, run, passage) and key genomic alterations (e.g., APC, TP53, KRAS/NRAS, FBXW7).

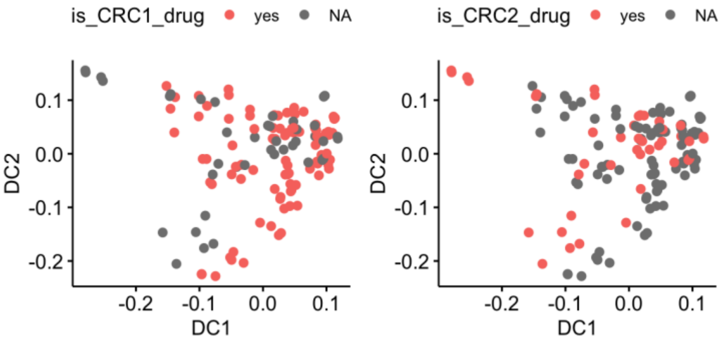


**Supp. Fig. 4. Distribution of two drug screening cohorts, CRC1 and CRC2, across the epithelial plasticity axes (DC1 and DC2).**

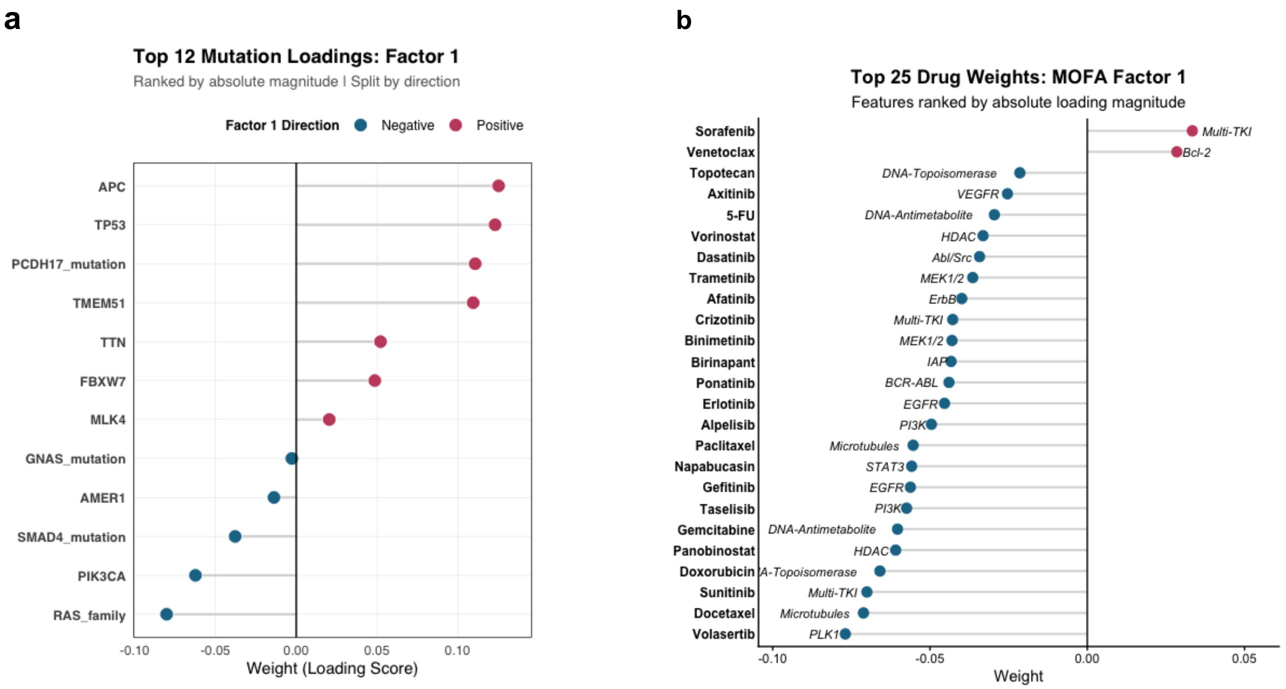


**Supp. Fig. 5. Gene and Drug loadings of MOFA Factor 1 (a)** Relative mutation loadings associated with latent factor 1 amounting for a total variability of 2.41%. (**b**) Top 25 drug loadings associated with factor 1.

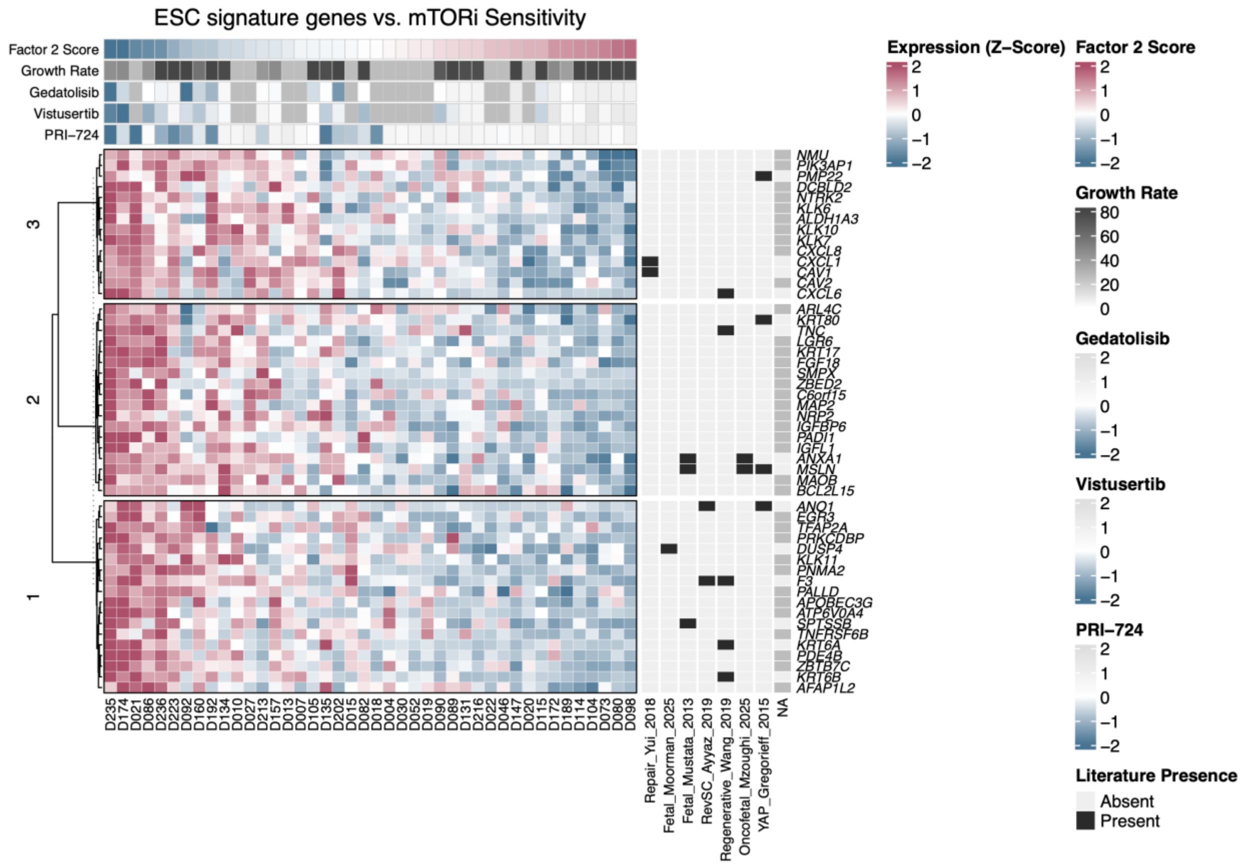


**Supp. Fig. 6. Transcriptional profile of embryonic and fetal-like stemness signature genes associated with FMD score.** Unsupervised hierarchical clustering of the top 50 candidate genes stratified into three distinct expression modules (Clusters 1–3). Columns represent individual patient-derived organoid (PDO) models (), ordered left-to-right by decreasing Factor 2 (FMD) score. Row values indicate row-wise Z-scaled normalized transcript expression counts, where red represents high expression and blue represents low expression. Top annotation denotes metadata mappings for each PDO, tracking the MOFA-computed loading for Factor 2 (FMD), the baseline proliferation rate, and drug sensitivity to Gedatolisib, Vistusertib & PRI-724 (Z-scored AUC). Right side panel contains uses binary annotation to highlight the presence or absence of FMD signature transcripts across landmark external published datasets characterizing intestinal regeneration, fetal stem cells and oncofetal states in colorectal cancer. Referenced source cohorts and markers in Supp. Table 3.

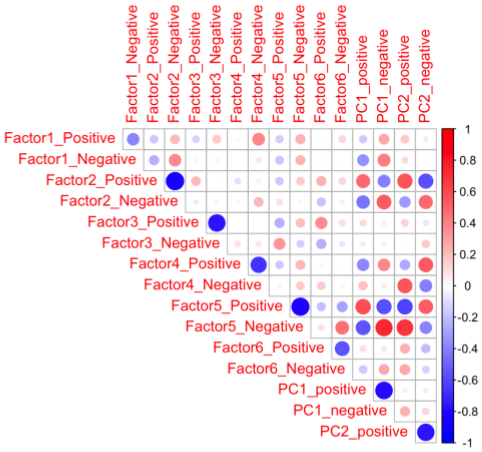


**Supp. Fig. 7. Correlation between MOFA factors and epithelial plasticity PCs.** Mean signature score is constructed by top 50 genes negatively or positively associated for PC and MOFA latent factors separately. Pearson correlation across each signature score is depicted from low (blue) to high (red). The sizes of the bubbles represent the magnitude (absolute value) of the correlation coefficient, with larger bubbles indicating stronger correlations and smaller bubbles indicating weaker correlations.

**a** **b**

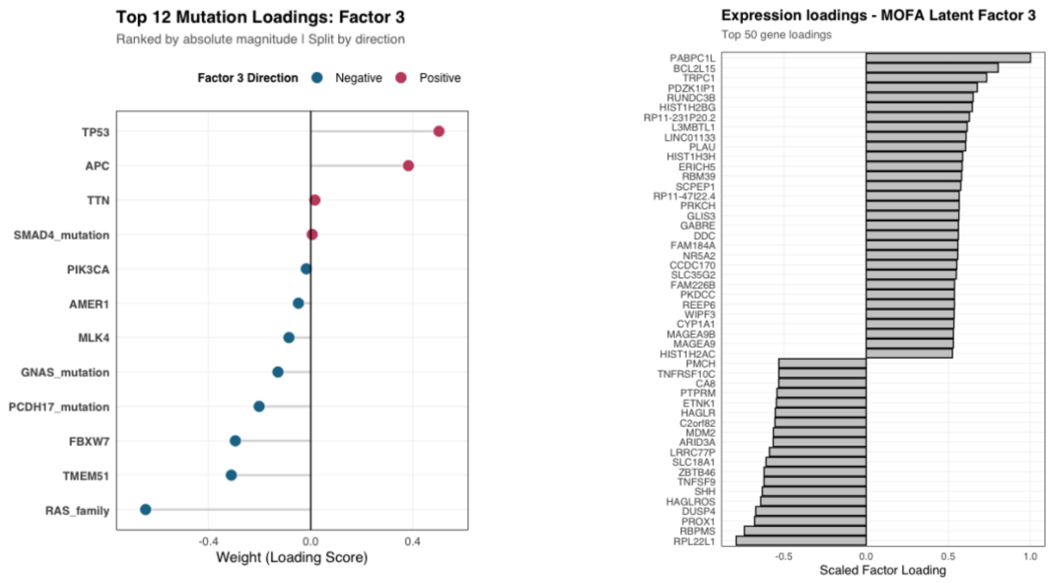


**Supp. Fig. 8. Mutation and gene loadings of Factor 3 (a)** Relative mutation loadings associated with latent factor 3 amounting for a total variability of 10.49%. **(b)** Top 50 gene loadings associated with latent factor 3

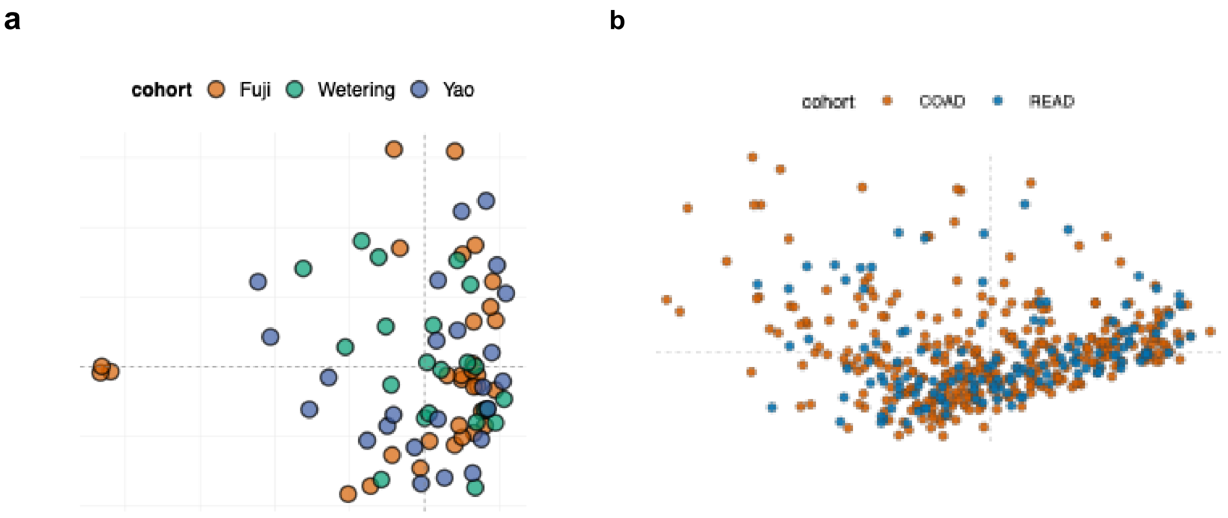


**Supp. Fig. 9. Distribution of different data sources across epithelial plasticity axes in (a) PDO biobanks and (b) TCGA CRC data.** Diffusion maps constructed from epithelial plasticity axis genes showing sample distributions. Samples are colored by data source, including study cohort in PDO biobanks and tumor site cohort in TCGA CRC data.

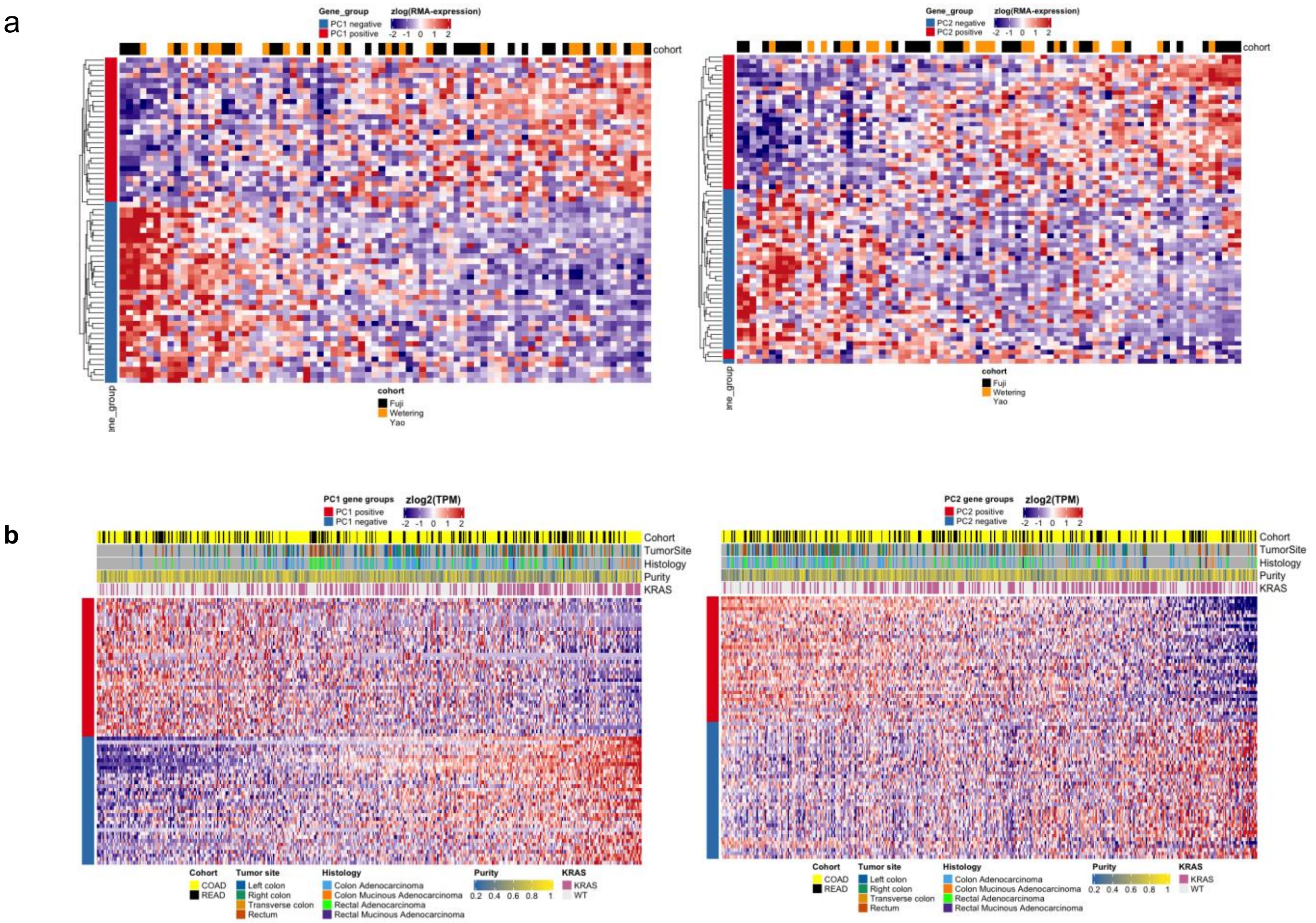
a
Gene_group
PC1 negative
PC1 positive
zlog(RMA-expression)
-2 -1 0 1 2
cohort
Gene_group
cohort
Fuji
Wetering
Yao
Gene_group
PC2 negative
PC2 positive
zlog(RMA-expression)
-2 -1 0 1 2
cohort
Gene_group
cohort
Fuji
Wetering
Yao
b
PC1 gene groups
PC1 positive
PC1 negative
zlog2(TPM)
-2 -1 0 1 2
Cohort
TumorSite
Histology
Purity
KRAS
Cohort
COAD
READ
Tumor site
Left colon
Right colon
Transverse colon
Rectum
Histology
Colon Adenocarcinoma
Colon Mucinous Adenocarcinoma
Rectal Adenocarcinoma
Rectal Mucinous Adenocarcinoma
Purity
0.2 0.4 0.6 0.8 1
KRAS
KRAS
WT
PC2 gene groups
PC2 positive
PC2 negative
zlog2(TPM)
-2 -1 0 1 2
Cohort
TumorSite
Histology
Purity
KRAS
Cohort
COAD
READ
Tumor site
Left colon
Right colon
Transverse colon
Rectum
Histology
Colon Adenocarcinoma
Colon Mucinous Adenocarcinoma
Rectal Adenocarcinoma
Rectal Mucinous Adenocarcinoma
Purity
0.2 0.4 0.6 0.8 1
KRAS
KRAS
WT

c

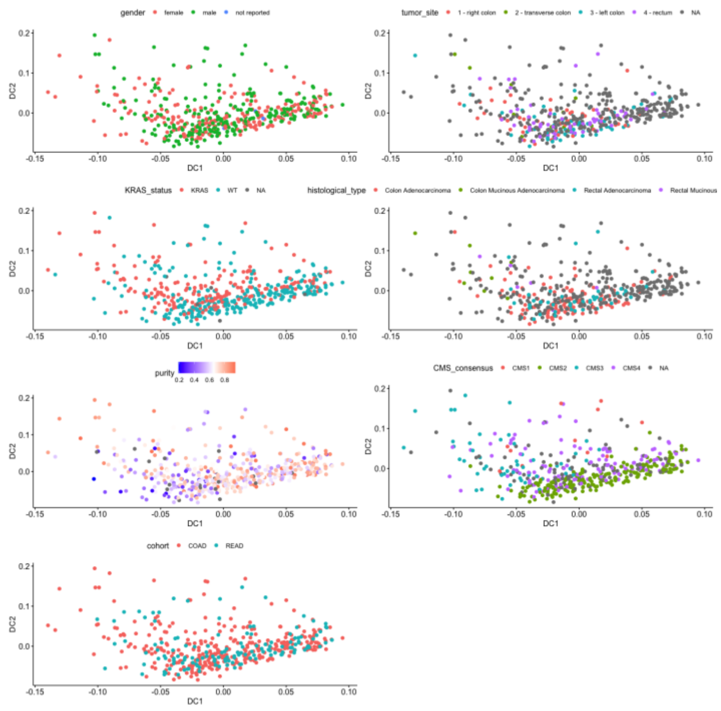


d

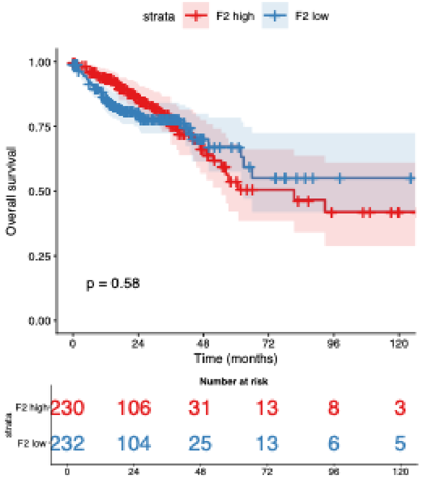


e

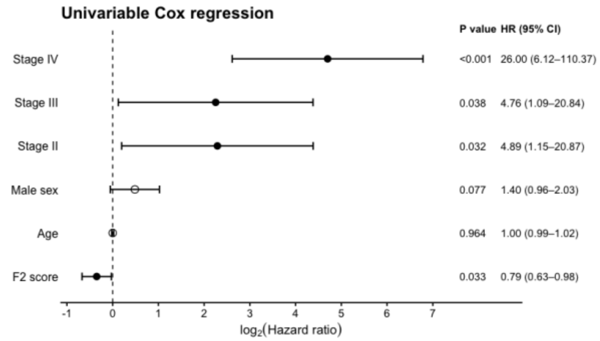


f

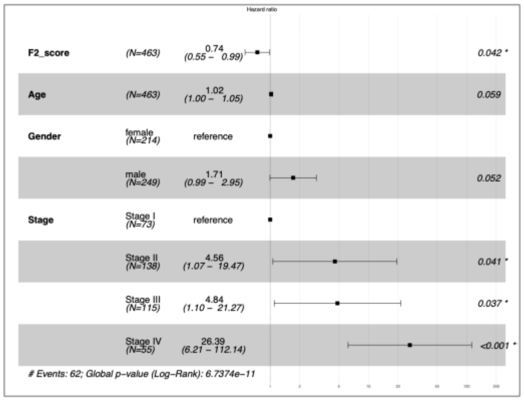


**Supp. Fig. 10. Validation of continuous epithelial axes across independent microarray and bulk RNA-seq CRC datasets.** Continuous expression of top 50 PC1 or PC2 genes across **(a)** three independent PDO biobanks and **(b)** TCGA CRC (COAD/READ). Z-scaled normalized expression values for each top PC1 (left) and PC2 (right) genes are depicted as heatmaps. Expression values are shown from low (blue) to high (red). Both sides of PC genes are colored separately: positive (red) and negative (blue) side. Samples are ordered along the DC1 from diffusion maps and genes are expressed along continuous gradients. **(c)** Clinical and genomic features on the epithelial plasticity axes in TCGA (COAD/READ). **(d)** Overall survival according to F2 score (FMD) in the TCGA CRC cohort. Patients are stratified into F2-high and F2-low groups based on the median F2 score. Kaplan–Meier survival curves are shown with 95% confidence intervals. P value was calculated using the log-rank test. **(e)** Univariable Cox regression analysis of F2 score (FMD) and clinical variables with respect to overall survival. Hazard ratios are shown on a log2 scale together with corresponding 95% confidence intervals (CI) and P values. **(f)** Multivariable Cox regression analysis including F2 score (FMD) as a continuous variable and adjusting for age, sex, and disease stage. Hazard ratios and 95% confidence intervals are shown, and P values are reported for each covariate.

a

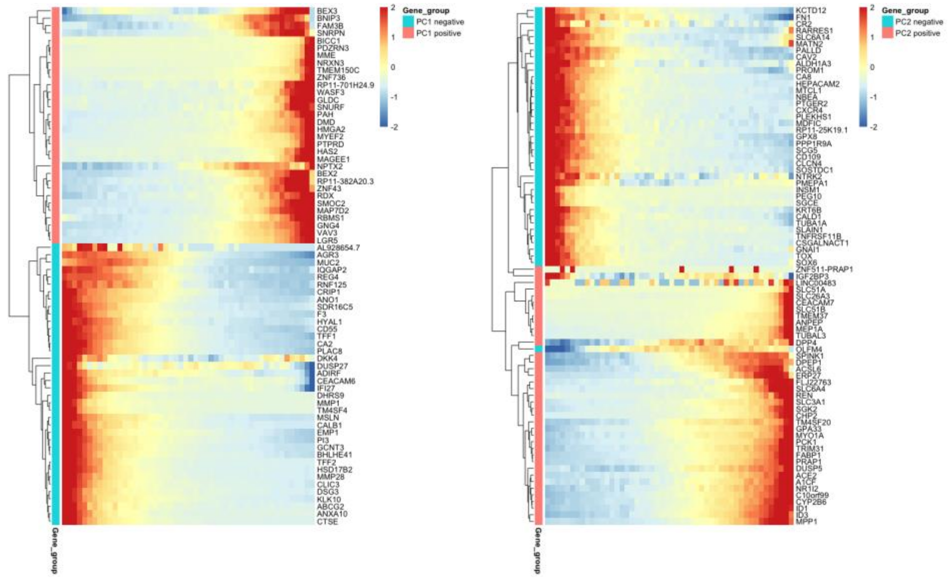

Gene_group
PC1 negative
PC1 positive
PC2 negative
PC2 positive
2
1
0
-1
-2
Gene_group


b

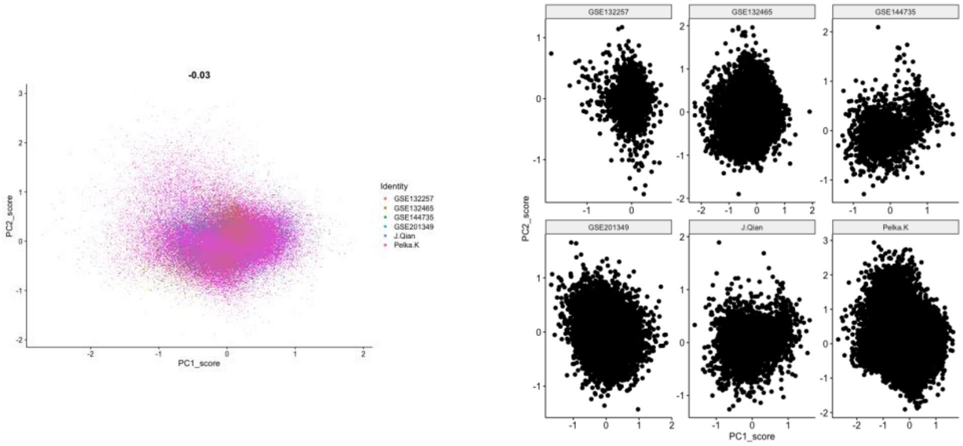

-0.03
Identity
GSE132257
GSE132465
GSE144735
GSE201349
J.Qian
Pelka.K
PC1_score
PC2_score
GSE132257
GSE132465
GSE144735
GSE201349
J.Qian
Pelka.K


c

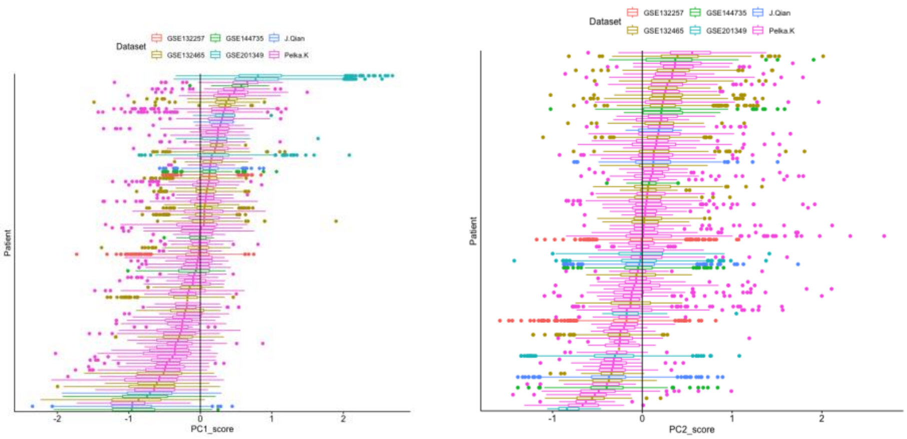

Dataset
GSE132257
GSE132465
GSE144735
GSE201349
J.Qian
Pelka.K
Patient
PC1_score
PC2_score

**Supp. Fig. 11. Validation of continuous epithelial axes across six independent single-cell CRC datasets. (a)** Continuous expression of top 50 PC1 or PC2 genes across single-cell CRC tumors. Malignant cells are used for the analysis. Cells are represented as bins of 50 cells each and Z-scaled normalized expression values for each top PC1 (left) and PC2 (right) genes are depicted as heatmaps. **(b)** Continuous representation of epithelial axes (PC1 and PC2). Signature scores are constructed for PC1 and PC2 epithelial programs by subtracting average expression of positive end genes from those from negative end. Datasets are colored and shown separately, left and right, respectively. Samples from datasets align across the epithelial programs. **(c)** Distribution of PC1 and PC2 scores across patients. PC1 and PC2 scores are shown separately as box plots for each patient, colored by datasets.

a

b

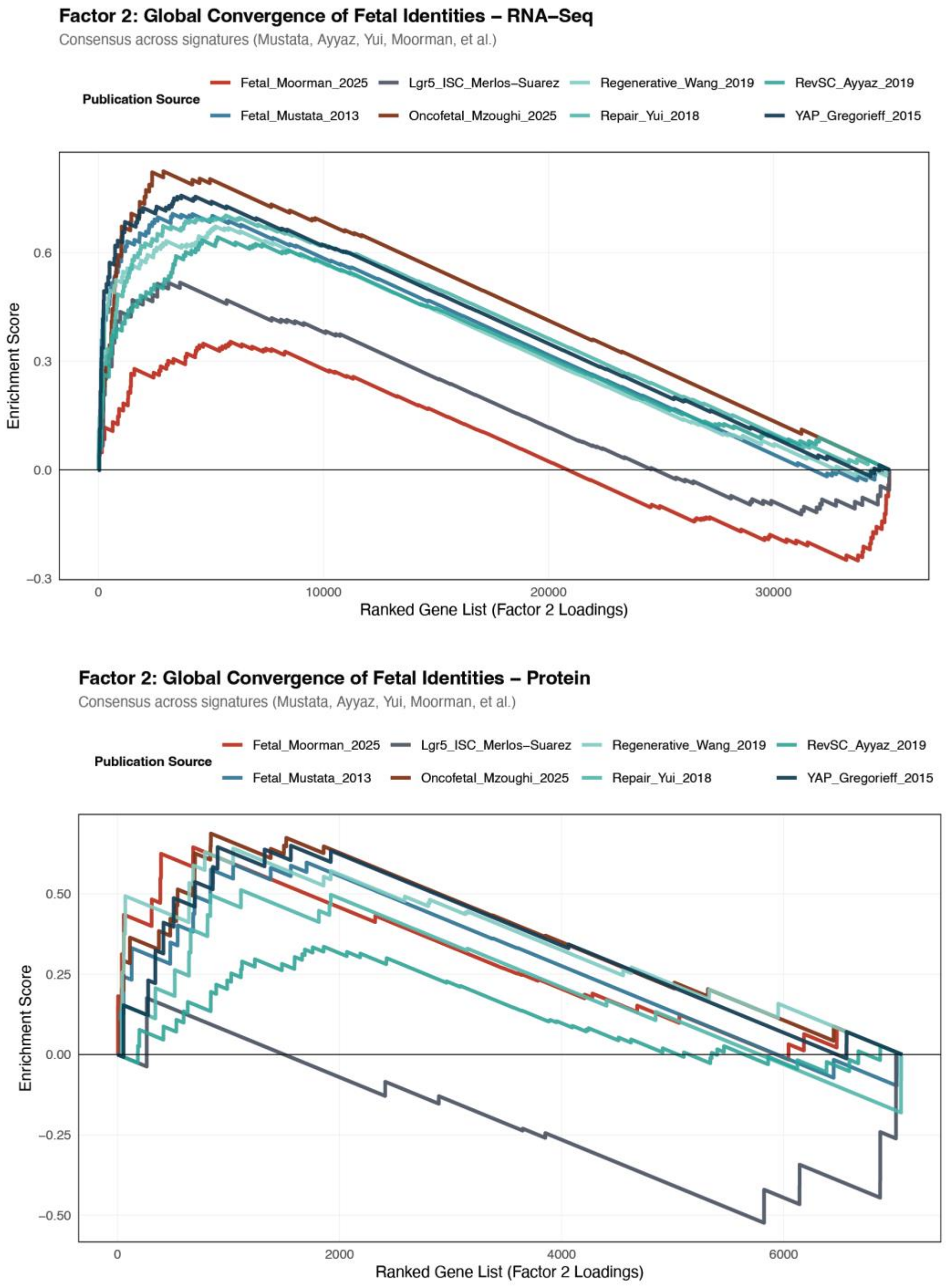


**Supp. Fig. 12. Multi-omic validation of fetal mTOR-dependency program (FMD).** Running enrichment score for fetal/regenerative intestinal stemness signatures ranked by differential expression between mTOR-sensitive and mTOR-resistant PDOs of validation cohort following RNA-Seq characterization **(a)** and mass spectrometry **(b)**
